# ArchVelo: Archetypal Velocity Modeling for Single-cell Multi-omic Trajectories

**DOI:** 10.1101/2025.09.14.676182

**Authors:** Maria Avdeeva, Sarah Walker, Joris van der Veeken, Alexander Rudensky, Yuri Pritykin

## Abstract

Inferring dynamic cellular processes from static single-cell measurements remains a central challenge in genomics. Here we introduce ArchVelo, a new method for modeling gene regulation and inferring cell trajectories using single-cell simultaneous chromatin accessibility (scATAC-seq) and transcriptomic (scRNA-seq) profiling. ArchVelo represents chromatin accessibility as a set of archetypes—shared regulatory programs—and models their dynamic influence on transcription. Compared to previous methods, ArchVelo improves inference accuracy and gene-level latent time alignment, and enables identification of the underlying transcription factor activity. We benchmark ArchVelo on developing mouse brain and human hematopoiesis datasets and apply it to CD8 T cells responding to viral infection, revealing distinct trajectories of differentiation and proliferation. Focusing on the progenitor CD8 T cell population with key roles in sustaining immune responses and translationally linked to immunotherapy outcomes, we identify a previously uncharacterized differentiation trajectory from Ccr6^−^ to Ccr6^+^ progenitors, shared between acute and chronic infection. In sum, ArchVelo provides a principled framework for modeling dynamic gene regulation in multi-omic single-cell data across biological systems.

## Introduction

Single-cell transcriptomics (scRNA-seq) has been widely used to characterize cellular heterogeneity across diverse biological systems.^1–5^ In scRNA-seq analysis, cells are represented as high-dimensional vectors of genome-wide gene expression, providing a quantitative multi-dimensional view of cellular phenotypes.^6–9^ These data are commonly structured as a *k*-nearest-neighbor (kNN) graph based on transcriptomic similarity between cells, which can be used to identify discrete cell types and states through clustering, or to reconstruct continuous biological processes such as cell differentiation or response to perturbations via trajectory inference.

Trajectory inference methods assume that cells undergo dynamic transcriptomic changes driven by underlying molecular processes, and that scRNA-seq provides snapshots of these processes across many cells. By analyzing the transcriptomic similarity graph, these methods aim to infer the progression of cellular states and assign pseudotime coordinates to cells. However, most trajectory inference methods operate on undirected graphs and rely on additional assumptions or external information to define origins, branch points, or endpoints of trajectories.^6, 10, 11^

RNA velocity methods represent a powerful class of trajectory inference approaches that exploit unspliced and spliced RNA reads in scRNA-seq data to model transcriptional dynamics directly.^12–20^ The ratio of unspliced to spliced reads provides a proxy for nascent versus mature transcript abundance, which can be modeled using ordinary differential equations (ODEs) to infer gene-specific regulatory kinetics. Aggregating these gene-level models allows estimation of a high-dimensional RNA velocity vector for each cell, enabling directional trajectory inference in gene expression space. While this framework has yielded valuable insights, it also presents challenges in terms of model robustness and interpretability.^14, 21^

Although gene expression reflects the output of regulatory processes, upstream regulatory activity, such as transcription factor binding and chromatin accessibility, can provide additional, potentially more direct, insights into cell state dynamics. In particular, scATAC-seq data, which measure chromatin accessibility, offer a window into regulatory element activity and can complement transcriptional data. Recent advances in multi-omic technologies now enable simultaneous profiling of chromatin accessibility and gene expression in the same cells (scATAC+RNA-seq), offering a more comprehensive view of gene regulation in dynamic cellular contexts.^4, 22–24^ However, modeling transcriptional dynamics from scATAC+RNA-seq data introduces additional complexity. Compared to scRNA-seq, scATAC-seq data are higher-dimensional, sparser, and lack readily available references of chromatin accessibility regions for preprocessing and quantification (unlike reference gene annotations for scRNA-seq analysis that are well curated).^6, 25^ Moreover, the relationship between chromatin accessibility and gene expression is not fully understood. We argue that a previously proposed method MultiVelo for RNA velocity analysis in scATAC+RNA-seq data^15^ could be improved by a more informative representation and modeling of scATAC-seq data.

Here we introduce ArchVelo, a new method for RNA velocity analysis and trajectory inference from scATAC+RNA-seq data. ArchVelo is based on archetypal analysis, which decomposes scATAC-seq profiles into a small number of interpretable archetypes representing “extreme” or characteristic regulatory programs within a dataset.^26, 27^ This low-rank representation of the chromatin accessibility modality improves RNA prediction performance and provides a biologically meaningful basis for modeling transcriptional dynamics. ArchVelo integrates this archetypal representation into a kinetic model that jointly describes the dynamics of chromatin accessibility and transcription. For each gene, transcriptional dynamics are modeled as a linear combination of contributions from the archetypes, enabling more accurate velocity estimation and trajectory inference. Using publicly available scATAC+RNA-seq data from developing mouse brain and human hematopoiesis, we show that ArchVelo achieves higher accuracy and improved latent time alignment for individual genes compared to state-of-the-art methods scVelo^13^ and MultiVelo.^15^ Furthermore, ground-truth benchmarking confirms that ArchVelo-inferred trajectories more closely recapitulate known biological progressions. Beyond improved accuracy, ArchVelo provides a novel feature: decomposition of the RNA velocity field into archetype-specific components. By coupling this decomposition with transcription factor (TF) motif analysis, ArchVelo enables mechanistic interpretation of cellular trajectories and nominates candidate TFs as drivers of specific regulatory components. To demonstrate the biological utility of ArchVelo, we applied it to a new multi-omic dataset profiling cytotoxic CD8 T cell responses to acute and chronic LCMV infection in mice. ArchVelo revealed two major orthogonal trajectories corresponding to functional differentiation and cell proliferation. Focusing on the Tcf1^+^ progenitor CD8 T cells, essential for sustained responses in cancer and infection and for cancer immunotherapies,^28–30^ we uncovered a previously uncharacterized differentiation trajectory from Ccr6^−^ to Ccr6^+^ progenitors.

In summary, ArchVelo is a robust and interpretable framework for RNA velocity analysis and trajectory inference from scATAC+RNA-seq data. By combining multi-omic modeling with archetypal decomposition, ArchVelo enhances the accuracy of dynamic inference and enables mechanistic insights into regulatory programs driving cell state transitions. The open-source ArchVelo package is available at https://github.com/pritykinlab/ArchVelo.

## Results

### Archetypal analysis of scATAC-seq data improves modeling of scRNA-seq data

A single-cell multi-omic sequencing experiment includes simultaneous genome-wide measurements of transcriptome (scRNA-seq) and chromatin accessibility (scATAC-seq) in single cells (**Fig. 1A**).

**Fig. 1.**
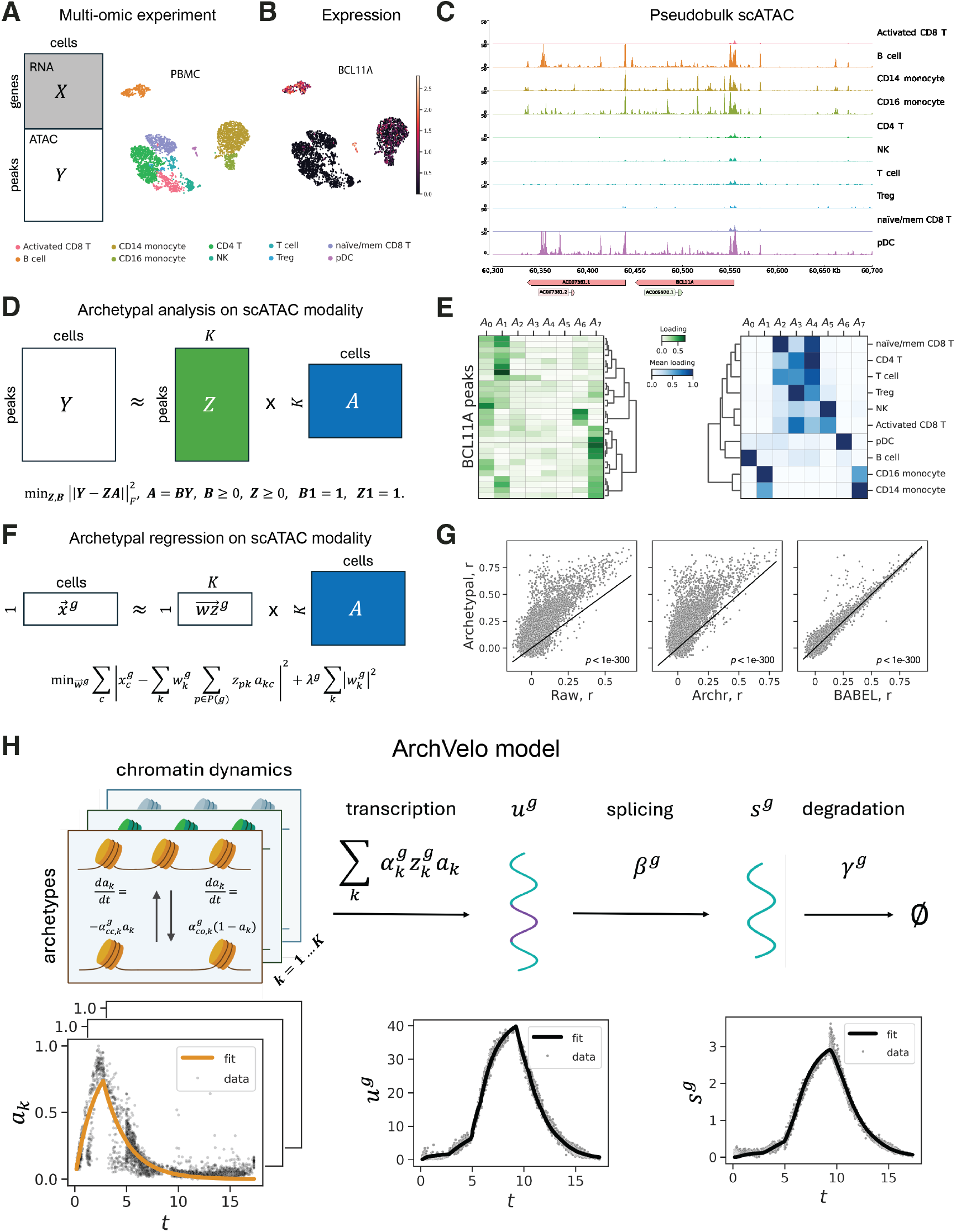
Single-cell ATAC-seq archetypes improve modeling of single-cell RNA-seq data. A multi-omic dataset of 12K human peripheral blood mononuclear cells (PBMCs) from 10x Genomics,^31^ subsampled to 3,000 cells, is used to illustrate archetypal analysis. **(A)** Left: A scATAC+RNA-seq experiment yields a gene-by-cell scRNA-seq matrix *X* and a peak-by-cell scATAC-seq matrix *Y*. Right: Uniform Manifold Approximation and Projection (UMAP) plot of the scRNA-seq data modality, with cell type annotations based on canonical marker genes. **(B)** Expression of gene BCL11A, which is active in multiple cell types. Normalized scRNA-seq expression. **(C)** Pseudo-bulk scATAC-seq signal at the BCL11A locus. **(D)** Schematic of archetypal analysis of the scATAC-seq data. After preprocessing and normalization, the chromatin accessibility profile of each peak is approximated as a convex combination of *K* archetypes (matrix *A*), which capture characteristic accessibility programs across the dataset. **(E)** Archetypal analysis with *K* = 8 applied to the PBMC dataset. Left: archetypal scores of peak summits (shorter, equally-sized regions of maximal accessibility within the peaks) in the BCL11A locus. Right: average cell type archetypal scores. **(F)** Schematic of scRNA-seq regression using scATAC-seq archetypes in scATAC+RNA-seq data. A ridge-regularized linear regression model is trained for each gene, using the archetypal accessibility features as predictors and scRNA-seq gene expression as the target. Cross-validation is used to select the regularization parameter. **(G)** Comparison of our regression model (panel F) with alternative approaches. Pearson correlation (*r*) between predicted and observed gene expression profiles in held-out test cells is plotted. Significance was assessed using the two-sided Wilcoxon signed-rank test. Left: baseline regression using raw scATAC-seq-derived peak accessibility linked to each gene. Center: ArchR^32^ gene activity scores. Right: predictions from BABEL.^33^ **(H)** Top: Schematic of the ArchVelo model. ArchVelo models the dynamics of chromatin accessibility, transcription, splicing, and degradation. Chromatin accessibility is represented by time-dependent profiles *a*_*k*_(*t*) which open and close over latent time *t*. Each archetype contributes linearly to the transcription rate of each gene, combining gene-level loadings 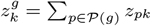 with archetype-specific transcription rates 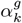. Bottom: Example model fits for a representative gene. Left: chromatin accessibility profile for one archetype *α_k_*(*t*) (orange line); middle: unspliced normalized imputed counts; right: spliced normalized imputed counts. Data points are shown in black.

The standard approach is to represent the scRNA-seq component as a matrix *X* of unique molecular identifier (UMI) or read counts for every gene in every cell, and to represent the scATAC-seq component as a matrix *Y* of read counts at peaks of chromatin accessibility in every cell. This scATAC-seq representation relies on a pre-constructed reference set of peaks, i.e. genomic regions with overall high level of scATAC-seq signal across cells. ATAC-seq peaks presumably correspond to sites of regulatory activity such as promoters, enhancers and repressors. Analysis of such data can be challenging due to high sparsity and dimensionality, and incomplete understanding of how regulatory activity, approximated with chromatin accessibility, drives gene expression. The chromatin accessibility landscape across cells can be quite complex, even within the locus of the same gene, where peaks may exhibit heterogeneous cell type-specific accessibility potentially associated with cell type-specific expression (**Fig. 1B,C**).

To address these challenges, we use archetypal analysis of the scATAC-seq data modality (**Fig. 1D,E**).^26, 27^ Each archetype *a*_*k*_ is a profile of chromatin accessibility over cells. For a pre-defined number of archetypes, every cell is decomposed into archetypal scores (contained in matrix *A*), and chromatin accessibility profile at every peak across cells is a convex combination of the archetypes (scores denoted by *z*_*pk*_ for peak *p*). To explore the global genome-wide relationship between the scATAC-seq and scRNA-seq modalities in a multi-omic dataset, we formulated a regularized linear regression model to predict scRNA-seq from scATAC-seq, using archetypal featurization of scATAC-seq (**Fig. 1F,G**). We observed that this simple approach had better predictive power than other methods. This includes scATAC-seq gene activity scores from ArchR, a popular package for scATAC-seq analysis,^32^ and a baseline regression approach using accessibility values in all peaks corresponding to a gene as features (**Methods**). The archetypal regression also performs significantly better than previously proposed deep learning method BABEL.^33^ Thus the low-dimensional archetypal representation of scATAC-seq data is productive for modeling the regulatory relationship between the two modalities in scATAC+RNA-seq data.

### ArchVelo for modeling dynamics of gene expression regulation

The standard analysis of scRNA-seq data uses estimates of mRNA abundance, i.e. the transcript product of gene expression. However, scATAC+RNA-seq analysis provides an opportunity to model the process of gene expression regulation rather than only its end product. Therefore, we next turned to modeling dynamics of gene expression regulation, extending the RNA velocity framework. The RNA velocity ordinary differential equations (ODEs)^12, 13^ model the cascade of transcription, splicing and degradation of individual genes assuming piecewise constant transcription rates, with different constant rates in induction and repression phases. This simplifying assumption was extended for scATAC+RNA-seq data in MultiVelo,^15^ where an additional equation modeling chromatin dynamics upstream of transcription was introduced, with the transcription rate proportional to the aggregated level of chromatin accessibility of a gene at any timepoint. However, this approach did not take full advantage of the resolution of the scATAC-seq data which measures chromatin accessibility at individual regulatory elements. Multiple regulatory programs can potentially drive transcription of a gene utilizing different subsets of its regulatory elements. To incorporate this information into the RNA velocity framework, we developed ArchVelo (**Fig. 1H**) that extends MultiVelo equations using the archetypal decomposition of the scATAC-seq modality (**Fig. 1D**). ArchVelo models dynamics of chromatin accessibility archetypes *a*_*k*_(*t*) for every gene *g* as a function of the underlying latent time *t*, incorporating individual transcription rates 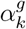 for every archetype, archetype accessibility score (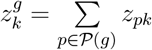 where 𝒫 (*g*) denotes the peaks mapping to *g*, **Methods**) and preserving the splicing and degradation model components *β*^*g*^ and *γ*^*g*^, respectively. The transcription rates are chosen to depend on transcription state 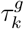 (**Methods**). The flexible kinetic model of ArchVelo (**Fig. 1H**), with dynamics constrained via joint scATAC-seq archetypal decomposition, could provide more accurate trajectory modeling than MultiVelo and give insight into dynamics of the archetypal regulatory programs driving transcription. To assess this and benchmark ArchVelo against previous methods, we next applied it to published and new datasets.

### ArchVelo enables improved trajectory inference in the mouse embryonic brain

Similar to other RNA velocity frameworks, ArchVelo combines gene-specific dynamic models to infer a global velocity field (**Methods**). To demonstrate its functionality and benchmark performance against other methods, we used a published scATAC+RNA-seq dataset of the embryonic mouse brain (**Fig. 2A**).^34^

**Fig. 2.**
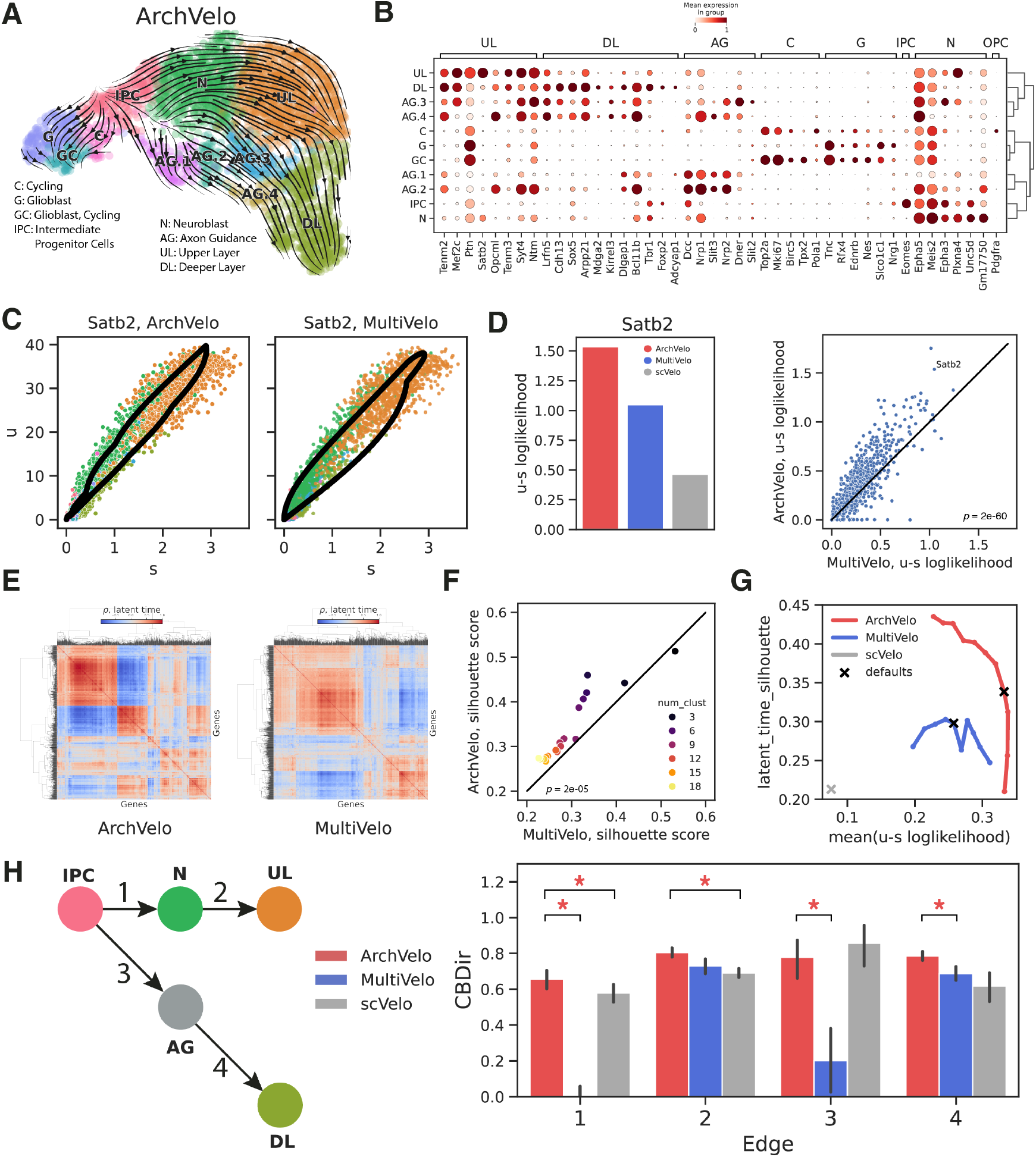
ArchVelo enables improved cell trajectory inference in the mouse embryonic brain. Comparison of ArchVelo with MultiVelo and scVelo on a multi-omic scATAC+RNA-seq dataset from mouse embryonic brain.^34^ **(A)** UMAP of the RNA modality with cell type annotations and stream plots showing velocity fields inferred by ArchVelo. **(B)** Dotplot of selected marker gene expression per cell type. Dot size reflects the proportion of expressing cells; color reflects min–max normalized expression. **(C)** Example velocity fits for gene *Satb2* in the unspliced-spliced (*u*–*s*) phase space, comparing ArchVelo and MultiVelo. **(D)** Left: log-likelihood (pseudocount = 1) of *u*–*s* phase space fits for *Satb2* using ArchVelo, MultiVelo, and scVelo. Right: scatterplot of per-gene log-likelihoods (*x*-axis: MultiVelo; *y*-axis: ArchVelo). Significance was assessed using the two-sided Wilcoxon signed-rank test. **(E)** Clustermaps of Spearman correlations between gene-specific latent time profiles inferred by ArchVelo and MultiVelo. **(F)** Scatterplot of silhouette scores for latent time correlation clusters (panel E) across varying cluster numbers. *x*-axis: MultiVelo; *y*-axis: ArchVelo. Significance was assessed using the two-sided Wilcoxon signed-rank test. **(G)** Benchmarking across all genes using two metrics: mean log-likelihood (pseudocount = 1) of *u*–*s* fits (*x*-axis) and silhouette score of latent time consistency (*y*-axis). Eleven values of the *w*_*c*_ parameter (ATAC-seq vs. RNA-seq contribution) were scanned for ArchVelo and MultiVelo. Default parameters are indicated. **(H)** Left: reference graph of known mouse embryonic brain differentiation. Nodes represent clusters (colored as in panel A; AG denotes the union of AG.1-4). Right: quantitative comparison of edge directionality using Cross-Boundary Direction (CBDir) scores^41^ for the reference graph edges. A star indicates significance for ArchVelo compared to another method (*p <* 0.01, Wilcoxon signed-rank test, Bonferroni corrected; only statistically significant comparisons shown).

We reanalyzed this dataset, identifying clusters corresponding to major developmental populations. These include glioblasts (G), intermediate progenitor cells (IPC), Unc5d^+^ neuroblasts (N), upper-layer (UL) and deeper-layer (DL) neurons. We also identified four neuronal subpopulations AG.1–AG.4 expressing the DL marker *Bcl11b* and various markers of axon guidance (**Fig. 2B, Fig. S1A**).^35^ In addition, we detected two distinct clusters of cycling cells: one enriched for glioblasts (GC) and another (C) comprising progenitors from both glial and neuronal lineages such as IPCs. ArchVelo revealed a separation of trajectories in the glial and neuronal compartment. Within the neuronal compartment, ArchVelo correctly inferred the separation of the upper and deeper layer lineages.^36, 37^ In particular, consistent with the literature, UL neurons were predicted to arise from IPC via Unc5d^+^ neuroblasts^38, 39^ while most DL neurons were inferred to originate from AG.1–AG.4, i.e., the Bcl11b^+^ neuroblasts or early neurons with strong expression of axon guidance signatures.^40^

We next compared the performance of ArchVelo with MultiVelo and scVelo on this dataset (**Fig. 2C–G, Fig. S1B,C**). As a result of its kinetic flexibility, ArchVelo produced more accurate fits of gene expression dynamics in the *u*–*s* phase space than MultiVelo (**Fig. 2C,D**). Both methods infer gene-specific latent times, but in ArchVelo, latent time estimation is constrained by shared chromatin archetypes, which improves consistency across genes. Consequently, ArchVelo produced better-aligned latent time profiles (**Fig. 2E,F, Methods**). Overall, owing to this shared structure and richer kinetic modeling, ArchVelo simultaneously achieved higher model likelihood and greater latent time robustness than MultiVelo across a wide range of the *w*_*c*_ parameters controlling ATAC-seq versus RNA-seq contributions (**Fig. 2G**). Cross-boundary direction correctness (CBDir) analysis^41^ confirmed that ArchVelo overall achieved higher scores than scVelo and Multi-Velo for biologically plausible transitions (**Fig. 2H, Methods**). Thus, ArchVelo more accurately recovered the differentiation dynamics in the developing brain than previous methods.

### ArchVelo recapitulates human hematopoietic stem cell differentiation trajectories and outperforms existing methods

We next applied ArchVelo to a multi-omic dataset of differentiating human hematopoietic stem cells (HSCs) previously generated in the MultiVelo study.^15^ In this experiment, purified CD34^+^ HSCs were cultured for seven days before sequencing. Our reanalysis revealed cell populations consistent with known hematopoietic differentiation hierarchies and the expression of lineage-specific markers (**Fig. 3A,B, Methods**).^42–48^

**Fig. 3.**
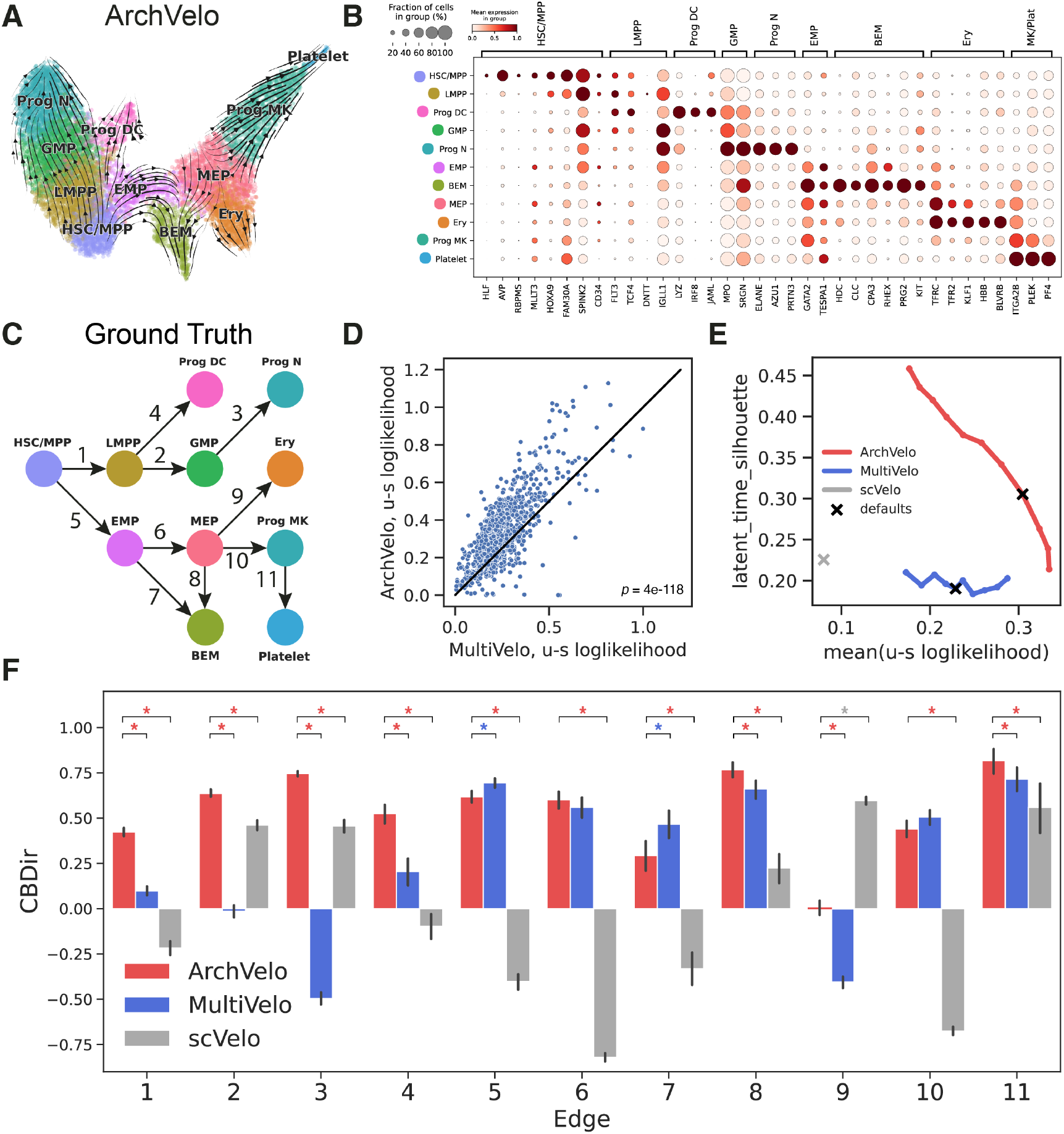
ArchVelo recapitulates human hematopoietic stem cell differentiation trajectories and outperforms existing methods. **(A)** UMAP of the HSC differentiation dataset with cell type annotations and velocity fields inferred by ArchVelo. HSC: human hematopoietic stem cells; MPP: multipotent progenitors; LMPP: lymphoid-primed multipotent progenitors; EMP: erythroid-myeloid progenitors; GMP: granulocyte–macrophage progenitors; Prog DC: progenitor dendritic cells; Prog N: progenitor neutrophils; BEM: basophil-eosinophil-mast progenitors; MEP: megakaryocyte–erythrocyte progenitors; Prog MK: progenitor megakaryocytes; Ery: early erythrocytes. **(B)** Dotplot showing expression of key cell type-specific marker genes across populations. Dot size indicates the proportion of expressing cells; color represents min–max normalized expression. **(C)** Reference graph of known hematopoietic differentiation (**Methods**). Nodes represent cell types (colored as in panels A and B). **(D)** Scatterplot comparing per-gene *u*–*s* log-likelihoods (pseudocount = 1) for MultiVelo (*x*-axis) and ArchVelo (*y*-axis). Significance was assessed using the two-sided Wilcoxon signed-rank test. **(E)** Benchmarking across all genes using two metrics: mean log-likelihood (pseudocount = 1) of *u*–*s* fits (*x*-axis) and silhouette score of latent time consistency (*y*-axis). Eleven values of the *w*_*c*_ parameter were scanned for ArchVelo and MultiVelo. Default parameters are indicated. **(F)** Quantitative comparison of edge directionality using Cross-Boundary Direction (CBDir) scores^41^ for the curated differentiation edges from panel C. Stars indicate significant differences (*p <* 0.01, Wilcoxon signed-rank test, Bonferroni corrected; only statistically significant comparisons shown); star color denotes the best-performing method.

In particular, we identified a subpopulation of HSCs closely resembling the multipotent progenitors (MPP), which seeded all major hematopoietic differentiation trajectories. Downstream differentiation branches included lymphoid-primed multipotent progenitors (LMPP), granulocyte–macrophage progenitors (GMP), progenitor neutrophils (N) and progenitor dendritic cells (DC). We also identified a subpopulation expressing intermediate levels of markers associated with erythroid (*TFRC, TFR2*), basophil-eosinophil-mast cell (*CPA3*), or both (*RHEX*) lineages. This mixed transcriptional profile, together with expression of *GATA2*, is consistent with erythroidmyeloid progenitors (EMP), a previously characterized early hematopoietic subpopulation.^45, 47, 48^ Additional clusters resembled megakaryocyte-erythroid progenitors (MEP), basophil-eosinophilmast cell progenitors (BEM), progenitor megakaryocytes (MK) and platelets, as well as early erythrocytes (Ery).

We constructed a curated reference graph of hematopoietic differentiation for the identified subpopulations (**Fig. 3C, Methods**).^44–48^ Notably, BEMs appear transcriptomically closer to erythroid and megakaryocyte progenitors than to other hematopoietic lineages.^43, 49^ Furthermore, early bifurcation between the monocyte–neutrophil–lymphocyte compartment and the megakaryocyte–erythroid–basophil–eosinophil–mast compartment has been proposed.^43, 45–47^ In the latter branch, since BEMs have been reported to arise from EMPs^45, 47^ and from MEPs,^43^ we incorporated both possibilities into the ground truth graph.

Evaluating trajectories inferred by ArchVelo against this curated reference, we found that ArchVelo successfully reconstructed most expected lineage relationships. Compared with MultiVelo, ArchVelo achieved higher per-gene log-likelihood (**Fig. 3D,E**) and greater consistency of latent time profiles (**Fig. 3E, Fig. S3A**). Cross-boundary direction correctness (CBDir) analysis^41^ further confirmed that ArchVelo achieved significantly higher or comparable CBDir scores than scVelo and MultiVelo (**Fig. S3B,C**) across nearly all lineage-defining edges of the reference differentiation graph (**Fig. 3F**). Unlike other methods, ArchVelo average CBDir scores were positive for all edges. The only edge with CBDir close to zero was the transition from MEP to Ery cells (edge 9) where MultiVelo predicted inverse directionality (**Fig. S3C**). Thus ArchVelo provides a more accurate and robust representation of hematopoietic differentiation dynamics than MultiVelo and scVelo.

Together these comprehensive benchmarking results demonstrate that ArchVelo outperforms previous methods in trajectory inference in single-cell multi-omic scATAC+RNA-seq data.

### ArchVelo enables decomposition of the velocity field into archetypal regulatory components

Beyond improving trajectory inference, ArchVelo provides a unique capability to decompose transcriptional dynamics into interpretable regulatory components (**Fig. 4A-C, Methods**). This is enabled by the linear structure of the underlying ODE model, where transcriptional variables *u*^*g*^(*t*) and *s*^*g*^(*t*) for each gene *g* can be expressed as linear combinations of archetype-specific contributions:

**Fig. 4.**
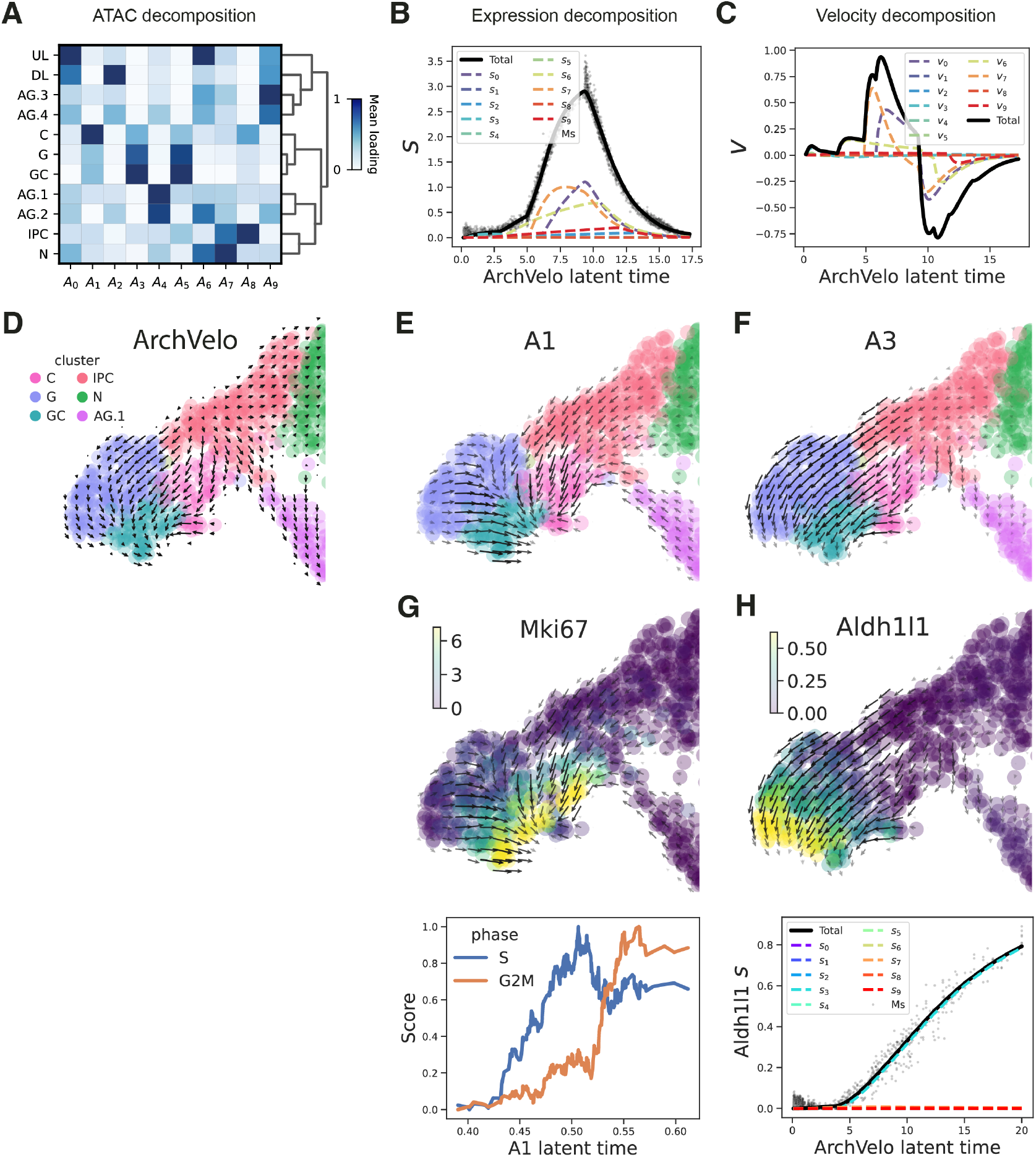
ArchVelo enables decomposition of the velocity field into archetypal regulatory components. Analysis was performed on the same multi-omic scATAC+RNA-seq dataset from embryonic mouse brain as in **Fig. 2**. **(A)** Clustermap of average archetypal loadings across annotated cell clusters. **(B-C)** Linear decomposition of ODE variables into components driven by individual chromatin accessibility archetypes. (B) Decomposition of spliced counts *s* for gene *Satb2* into components *s*_*k*_ associated with archetypes *a*_*k*_. Colored dashed lines show individual components; the black line indicates the total predicted *s*. The scatterplot shows imputed spliced counts. **(C)** Same decomposition for the velocity 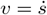, using the same color scheme as in panel B. **(D)** Grid visualization of the ArchVelo velocity field in the UMAP, focusing on the glial compartment. **(E-F)** Example velocity components for archetypes A1 (E) and A3 (F), shown on the same grid as in panel D. While both components are concentrated on similar cell subsets, they predict distinct local velocity patterns, reflecting cell cycle progression (A1) and astrocyte lineage differentiation (A3), respectively. **(G)** Top: velocity field for A1 overlaid with imputed expression of the proliferation marker *Mki67*. Bottom: cell cycle phase scores (S and G2M), inferred from gene signatures, plotted against latent time estimated from A1 (smoothed over 100 nearest neighbors), demonstrating a strong correspondence with cell cycle dynamics. **(H)** Top: velocity field for A3 overlaid with imputed expression of *Aldh1l1*, a canonical astrocyte marker that initiates in radial glia and is progressively upregulated during astrocyte lineage differentiation. Bottom: decomposition of spliced counts for *Aldh1l1* into archetypal components, as shown in panel B.

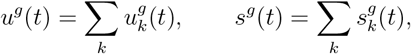

where 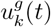 and 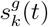 denote the components driven by archetype *a*_*k*_(*t*). Consequently, the gene-specific velocity

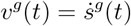

is similarly decomposable into archetypal components:

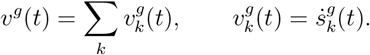

Each component represents the portion of a gene’s expression and velocity attributable to regulatory elements associated with a particular chromatin accessibility archetype (**Fig. 4A-C**).

Aggregating these contributions across all genes enables ArchVelo to extract distinct components of the global velocity field, each potentially corresponding to a specific regulatory program (**Methods**). This decomposition is computed at the single-cell level, and the resulting components can be directly projected onto the UMAP embedding, revealing archetype-specific trajectories that can be independently visualized and interpreted.

For example, in the embryonic mouse brain dataset, two archetypes (A1 and A3) produced velocity fields concentrated in the glioblast compartment (**Fig. 4D-H, Fig. S2A,C**). Although they overlapped in cell coverage, the inferred dynamics were clearly distinct. Archetype A1 was enriched in cycling cells and associated with cell cycle-related genes such as *Top2a* and *Kif23* (**Fig. S2B**). Consistent with this, the latent time inferred from A1 aligned with progression through cell cycle phases (**Fig. 4G**). In contrast, archetype A3 encompassed both cycling and non-cycling glioblasts and was enriched for astrocyte lineage markers, including *Aldh1l1* and *Slc1a3* (**Fig. 4H, Fig. S2D**). The corresponding velocity field suggested differentiation trajectories toward astrocyte progenitors, supported by the progressive upregulation of *Aldh1l1* during astrocyte lineage commitment. Thus, ArchVelo disentangled proliferation and differentiation components within the same precursor population, clarifying trajectories of astrocyte-committed glioblasts.

These findings highlight how ArchVelo’s archetypal decomposition disentangles the superposition of multiple regulatory influences that simultaneously shape the trajectory of a cell. In conventional trajectory analyses, the observed dynamics represent an aggregate view of all underlying forces, which can obscure distinct processes such as cell cycle progression and lineage commitment. By resolving the velocity field into archetype-specific components, ArchVelo isolates these regulatory contributions, enabling separate interpretation of concurrent programs and clarifying how different forces drive cells in potentially divergent directions.

In the HSC dataset, decomposition of the velocity field highlighted archetypes A0, A2, and A3, which revealed diverging trajectories from the progenitor MEP subpopulation toward megakaryocyte (Prog MK), BEM and erythroid (Ery) progenitors, respectively (**Fig. 5A,B**). TF motif analysis for these archetypes uncovered top factors potentially driving these differentiation trajectories (**Fig. 5C–E, Methods**). For example, the top motif for archetype A0 corresponded to factors ERG, ETV3, and FLI1, with FLI1 being a well-established driver of megakaryocyte identity.^50^ Consistent with this role, FLI1 motif scores, expression, and the expression of its downstream target ITGA2B showed expected patterns (**Fig. 5C**).^51^ Analogously, GATA2 emerged as the top regulator of BEM cell differentiation (A2, **Fig. 5D**), while the GATA family and KLF1 scored highest for the erythroid trajectory (A3, **Fig. 5E**), consistent with their well-established roles in these lineage specifications.^52, 53^ Together, these results validate the ability of ArchVelo decomposition to link distinct velocity components to candidate TFs driving lineage-specific regulatory programs. Moreover, as illustrated in the hematopoiesis analysis, individual archetypal components can in some cases correspond to *bona fide* lineage bifurcations, rather than merely overlapping regulatory influences, thereby directly uncovering the branching structure of differentiation trajectories.

**Fig. 5.**
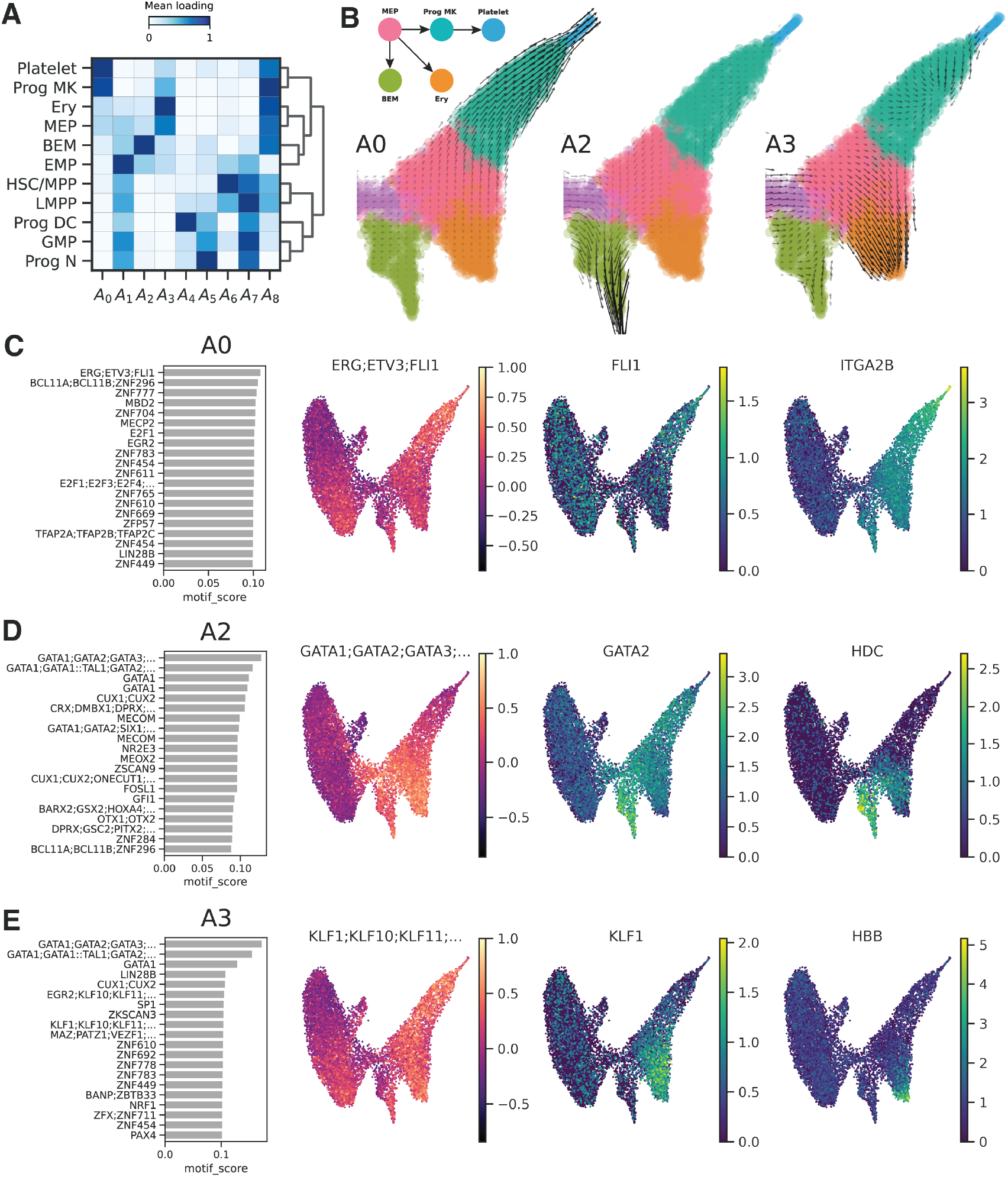
ArchVelo identifies candidate transcription factor drivers of distinct regulatory components of RNA velocity through motif analysis. Analysis was performed on the same multi-omic scATAC+RNA-seq dataset of human HSCs as in **Fig. 3**. **(A)** Heatmap showing average archetypal loadings across annotated cell clusters. **(B)** Velocity field components corresponding to archetypes A0, A2, and A3, visualized on a common embedding grid (same UMAP as in **Fig. 3A**). Each component captures a distinct biologically plausible differentiation trajectory. **(C)** Transcription factor (TF) motif analysis for archetype A0. Left: Barplot of top-scoring motifs for A0 (**Methods**). Right: Per-cell TF motif scores, normalized scRNA-seq expression of a representative gene encoding the corresponding TF, and a marker gene that is a downstream target of the TF. **(D)** Same as panel C, for archetype A2. **(E)** Same as panel C, for archetype A3.

In sum, decomposition of ArchVelo trajectories into multiple, potentially divergent trajectories driven by different regulatory programs within the same cells, represented by scATAC-seq archetypes, provides a conceptually new view of regulatory dynamics beyond cell trajectories.

### ArchVelo identifies distinct trajectories of cell differentiation and proliferation in CD8 T cells responding to viral infection

We next applied ArchVelo to a distinct biological system, cytotoxic CD8 T cells responding to viral infection. Specifically, we analyzed a multi-omic scATAC+RNA-seq dataset collected at day 7 post-infection with either the Armstrong (acute) or clone 13 (chronic) strains of lymphocytic choriomeningitis virus (LCMV). This dataset was generated as part of our larger recent study characterizing splenic T cell populations across infection conditions, building on our prior work.^54–56^

Using scATAC-seq accessibility signatures and gene activity profiles, we first identified antigen-experienced CD8 T cells, the population most relevant for studying infection-induced dynamics (**Methods**).^54^ Within this compartment, we refined cell state annotations, resolving multiple sub-populations consistently observed in both LCMV Armstrong and clone 13 infections (**Fig. 6A– C**). Marker gene expression confirmed activated effector (e.g., *Klrg1, Ccl5, Gzmb, Gzmk, Gzma*), memory-like effector (e.g., *Il7r*), and exhausted (e.g., *Pdcd1, Tox*) states, as well as intermediate subpopulations, alongside distinct cycling subsets characterized by *Mki67, Top2a, Pola1* and *Pcna* (**Fig. 6A,B**). Cycling cells were largely devoid of terminal differentiation markers, suggesting independent regulatory programs. We also identified a progenitor population defined by expression of *Tcf7* (encoding protein Tcf1), *Cxcr5, Slamf6, Id3, Sell* and *Il7r* but lacking activation-associated genes *Gzmb, Ccl5* and *Prf1*. This well-studied subset has been implicated in long-term immune responses in infection and cancer and linked to immunotherapy outcomes.^28–30^ Consistent with recent work by us and others, we found the progenitors present in both acute and chronic infection (**Fig. 6A–C**).^56–59^ Separately, we resolved an effector subpopulation with high expression of activation genes (*Gzmb, Ccl5, Prf1*) but also enriched for the memory marker *Il7r*, approaching levels seen in progenitors. Importantly, all identified subsets were shared between acute and chronic responses (**Fig. 6A–C**). This refined annotation provided the foundation for applying ArchVelo to dissect dynamic trajectories in infection-induced CD8 T cell responses.

**Fig. 6.**
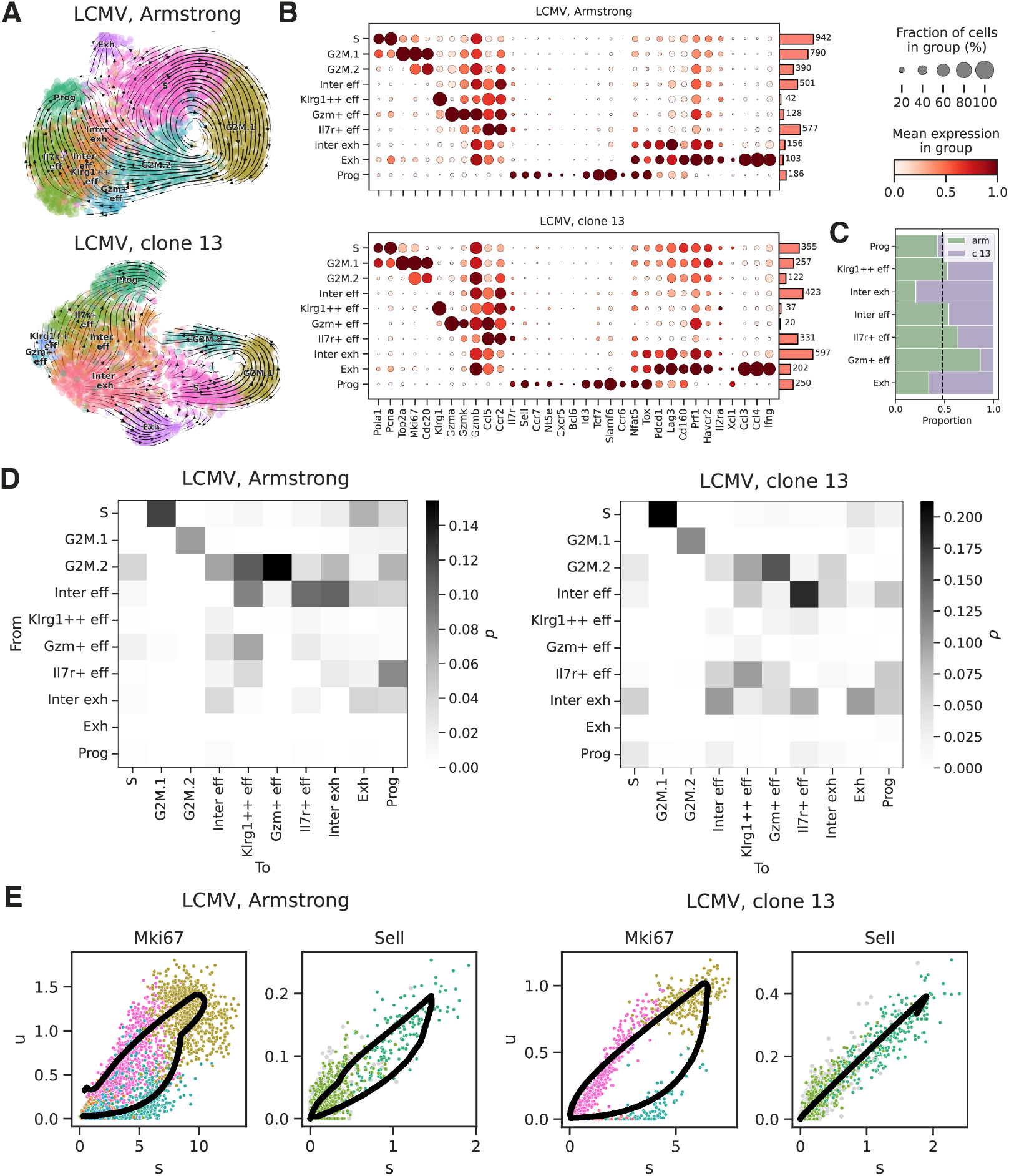
ArchVelo uncovers shared trajectories of cell differentiation and proliferation in CD8 T cells responding to acute and chronic viral infection. ArchVelo analysis of a multi-omic scATAC+RNA-seq dataset for cytotoxic CD8 T cells at day 7 post-infection with either acute (Armstrong) or chronic (clone 13) lymphocytic choriomeningitis virus (LCMV). **(A)** UMAP visualization of the scRNA-seq modality for Armstrong (top) and clone 13 (bottom), overlaid with ArchVelo stream plots. Datasets were clustered jointly. Cluster annotations: S: S phase; G2M.1: early G2/M phase; G2M.2: late G2/M phase; Inter eff: intermediate effector cells; Klrg1^++^ eff: effector cells expressing the highest levels of *Klrg1*; Gzm^+^ eff: *Gzma*- and *Gzmk*-expressing effector cells; Il7r^+^ eff: *Il7r*-expressing effector cells; Inter exh: intermediate exhausted cells; Exh: exhausted cells; Prog: progenitor cells. **(B)** Dotplot of selected marker gene expression across annotated clusters, shown separately for each LCMV clone. Dot size reflects the proportion of expressing cells; color indicates min–max normalized expression; bar height shows cluster size. **(C)** Clonal composition of non-cycling clusters. Dashed vertical line: expected proportion based on clone-specific total non-cycling cell numbers. **(D)** Heatmaps of ArchVelo transition probabilities between clusters (**Methods**), for the two infection conditions. **(E)** Phase plots for genes *Mki67* and *Sell* for the two infection conditions. Cluster colors are the same as in panel A. Black curve: ArchVelo fit.

ArchVelo revealed two major largely orthogonal sets of trajectories: one corresponding to cell cycle progression and the other to functional T cell differentiation (**Fig. 6A,D–F**). Transitions between cell states were consistent across both infection clones and could be further illustrated by phase plots of ArchVelo fits for individual genes (**Fig. 6D,E, Fig. S4A,B, Methods**). For example, *Mki67* dynamics aligned with transitions through cell cycle phases, while *Sell* expression captured differentiation from Il7r^+^ effectors into progenitors. In sum, ArchVelo identified distinct trajectories of CD8 T cell differentiation and proliferation that were largely shared between responses to LCMV Armstrong and clone 13.

To further leverage the resolution of ArchVelo, we focused on the CD8 T cell progenitor compartment, a population of major translational interest. Within this compartment, we identified two subpopulations of Ccr6^−^ and Ccr6^+^ progenitor CD8 T cells, present in both Armstrong and clone 13 infections (**Fig. 7A,B**). Both subsets expressed canonical progenitor markers, but also exhibited distinct gene expression profiles, indicating functional heterogeneity. This observation is consistent with previous reports of analogous Ccr6^−^ vs. Ccr6^+^ progenitor subsets in cancer, as well as CD62L^+^ vs. CD62L^−^ progenitors in chronic infection, whose gene signatures resembled those of our Ccr6^−^ and Ccr6^+^ subsets, respectively (**Fig. S5A**).^60–62^ In cancer, the Ccr6^+^ subset was reported to respond poorly to checkpoint blockade, with functional responsiveness instead attributed to the less differentiated Ccr6^−^ subset.^60^ Consistent with this, in LCMV infection we observe that Ccr6^−^ progenitors differentiate to the Ccr6^+^ progenitors (**Fig. 7A,B, Fig. S5B**).

**Fig. 7.**
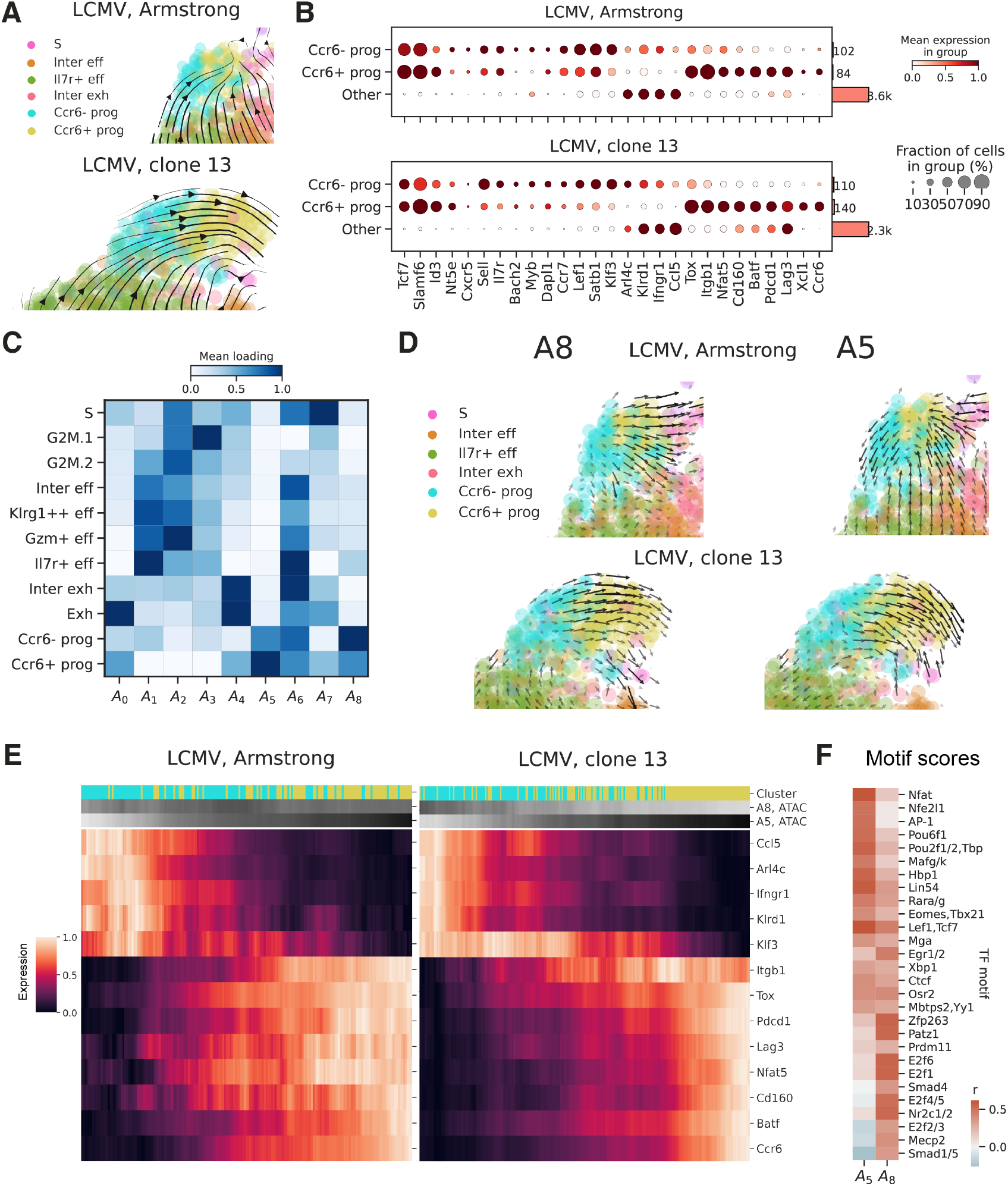
ArchVelo uncovers a differentiation trajectory from Ccr6^−^ to Ccr6^+^ progenitor CD8 T cells. The analysis focused on the progenitor compartment of cytotoxic CD8 T cells at day 7 following acute (Armstrong) or chronic (clone 13) LCMV infection (same data as in **Fig. 6**). **(A)** Zoomed-in view of ArchVelo-inferred trajectories within the progenitor CD8 T cell compartment and adjacent clusters in Armstrong (top) and clone 13 (bottom). The Ccr6^−^ and Ccr6^+^ progenitor subpopulations are highlighted. **(B)** Dotplots showing expression of known progenitor marker genes across Ccr6^−^ progenitors, Ccr6^+^ progenitors, and all other cells, as well as genes differentially expressed between Ccr6^−^ and Ccr6^+^ progenitors. Dot size indicates the proportion of expressing cells; color indicates min–max normalized expression; bar height represents cluster size. **(C)** Heatmap of average archetypal loadings across annotated clusters, incorporating the refined annotation of progenitor subpopulations. **(D)** Velocity field components corresponding to archetypes A8 (left) and A5 (right), visualized on the same UMAP embedding as in **Fig. 6A**. Top: Armstrong; bottom: clone 13. **(E)** Archetypal decomposition reveals a shared differentiation trajectory across clones within the progenitor subpopulation. Heatmaps show min–max normalized imputed scRNA-seq expression of selected genes (averaged over a sliding window of 20 cells), aligned by latent time. Top color bar: Ccr6^−^ and Ccr6^+^ progenitor annotations. Middle and bottom color bars: min–max normalized archetypal loadings for archetypes A8 and A5, respectively (averaged over a sliding window of 20 cells). Differentially expressed genes from panel B are shown if ArchVelo models were fit successfully. **(F)** Heatmap of TF motif scores for archetypes A5 and A8 within the progenitor subpopulation. Top 5 motives for every archetype are included.

Archetypal loadings reinforced the distinction between the Ccr6^−^ and Ccr6^+^ progenitor subsets and highlighted their associated velocity components A8 and A5, respectively (**Fig. 7C,D**). Archetype A8 was associated with the transition from Ccr6^−^ to Ccr6^+^ progenitors, whereas archetype A5 appeared to capture further differentiation of Ccr6^+^ progenitors and was weaker in the Ccr6^−^ progenitors. Using archetypal latent times, we reconstructed a shared differentiation trajectory across both infection contexts (**Fig. 7E**). This analysis revealed smooth and coherent gene expression dynamics, encompassing many of the differentially expressed genes (**Fig. 7B**), and supported a continuous progression between progenitor subsets. TF motif analysis of these archetypal components identified candidate factors driving this progression (**Fig. 7F**). In particular, archetype A8, associated with the less differentiated Ccr6^−^ progenitors, was enriched for Smad and E2f family motifs, consistent with regulatory programs maintaining proliferative potential. In contrast, archetype A5, linked to further differentiation of Ccr6^+^ progenitors, showed enrichment of Nfat and AP-1 family motifs, factors with well-established roles in T cell activation and effector differentiation. Thus using ArchVelo we identified a previously undescribed differentiation from Ccr6^−^ to Ccr6^+^ progenitor CD8 T cells in both acute and chronic LCMV infection.

Together, these results demonstrate that ArchVelo enables fine-grained dissection of overlapping dynamic programs such as proliferation and differentiation, both hallmarks of active T cell response *in vivo* to antigen stimulation. By uncovering shared trajectories across infection states, ArchVelo offers a powerful framework for understanding cellular decision-making.

## Discussion

We presented a new method ArchVelo for improved modeling of gene expression regulation dynamics and trajectory inference using multi-omic scATAC+RNA-seq data. Methodologically, two elements were critical. First, representing scATAC-seq as archetypes mitigates sparsity and shares information across regulatory elements with similar cell-wise activity, yielding stronger RNA prediction and a more stable regulatory basis for kinetic modeling. Second, embedding this representation into a gene-wise ODE system lets each archetype contribute linearly to transcription, providing flexible kinetics without sacrificing interpretability. Together, these choices improved quantitative performance (higher *u*–*s* likelihoods; more coherent latent times) and produced velocity fields that better captured expected developmental progressions. We applied ArchVelo across three biological systems, embryonic mouse brain, human hematopoiesis, and virus-responding CD8 T cells, recovering expected lineage progressions in the first two and, in T cells, delineating orthogonal proliferation and differentiation programs and a previously uncharacterized trajectory from Ccr6^−^ to Ccr6^+^ progenitors across acute and chronic infection.

We also combined ArchVelo archetypal trajectory decomposition with TF motif analysis, enabling nomination of candidate TFs associated with distinct regulatory components. While these associations are correlative, they provide testable hypotheses about TF programs that couple chromatin dynamics to transcriptional change and may guide perturbation studies. These results highlight the potential of integrating regulatory analysis with velocity modeling. In future work, ArchVelo could be extended with a more seamless framework to systematically connect TF activity, chromatin accessibility dynamics, and transcriptional trajectories in multi-omic data.

ArchVelo is a new method in the broader family of trajectory inference methods built on the RNA velocity framework. Many such approaches proceed in two stages: first, fitting a dynamic model for each gene independently by leveraging variation across cells; second, aggregating these gene-level fits into high-dimensional velocity fields that are then combined into trajectories. In ArchVelo, archetypal representation of scATAC-seq data improves both stages by stabilizing gene-wise modeling and producing more coherent velocity fields. Looking forward, this concept could be extended further by merging the two stages into a joint model of gene regulation across many genes simultaneously, moving beyond purely gene-by-gene modeling. The archetypal decomposition of scATAC-seq already provides a natural foundation for this direction.

Thus, ArchVelo illustrates a productive new direction for RNA velocity modeling, combining improved accuracy with interpretability. Beyond its immediate applications, the framework provides a foundation that can be extended in multiple ways, opening opportunities for future methodological advances in single-cell multi-omics.

## Methods

### Archetypal analysis on ATAC modality

We reduce dimensionality of the ATAC modality of each multi-omic dataset by applying archetypal analysis (AA).^26, 27^ For an experiment with *N* cells and *S* peak summits, let us denote the *S* × *N* Pearson residual normalized matrix of peak or peak summit accessibility profiles as *Y*. For a fixed dimensionality parameter *K*, AA solves the following optimization problem:

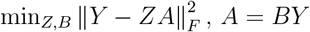

where *B* and *Z* satisfy additional constraints *B* ≥ 0, *Z* ≥ 0, *B*1 = 1, *Z*1 = 1. The matrix *A* ∈ ℝ^*K×N*^ contains the archetypal chromatin accessibility profiles in the rows. The condition *Z*1 = 1 on the *S* × *K* matrix of non-negative peak summit loadings ensures that the accessibility profile of every summit, i.e., every row of the ATAC matrix *Y*, is approximated by a convex combination of archetypes. Moreover, the conditions on the *K* × *S* matrix *B* ensure that every archetype lies on the boundary of the convex hull of the accessibility profiles at peak summits. AA can be viewed as a soft clustering technique without the orthogonality constraint. We solve the optimization problem using the efficient implementation of the AA algorithm by Mørup et al^63^ available via py_pcha Python package. The implementation was developed for large scale AA, and is based on projected gradient as well as an efficient initialization procedure. The implementation also includes a relaxation of the AA problem where the archetypes can reside outside of the convex hull of the observations which the authors call the AA-*δ* problem. In our notation, the problem modification relaxes the *B*1 = 1 condition into max|*B*1 − 1|*< δ*. By default, we use *δ* = 0.1. For more details on archetypal analysis and the AA-*δ* algorithm we refer the reader to available literature.^26, 27, 63^

### Linear model to predict RNA from ATAC modality

We train a model to predict the log-transformed library-size normalized RNA count matrix *X* from the ATAC modality *Y*. We choose to use the archetypes as features in a ridge-regularized linear regression model:

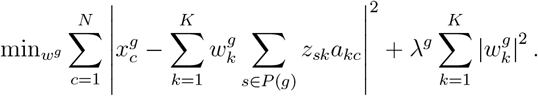

Here *c* and *k* are the cell and archetype indices, *g* denotes a gene and its accessibility score for archetype *k* is given by the sum of the loadings *z* over the set of all peak summits *P* (*g*) mapping to *g*. We train the model for every gene independently. We assume a linear relationship between the number of gene transcripts and archetypal chromatin accessibility of the gene, with the desired coefficients 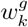 describing the overall input of every archetype into the pool of transcripts.

For training and testing purposes we separate every dataset into independent subsets, using *N*_train_ cells for *Y*_train_ and *N*_test_ cells for *Y*_test_. By default, we use 75% and 25% of the data for these purposes, respectively. To train and test our linear model, we first apply AA-*δ* on *Y*_train_ to extract the archetypal chromatin accessibility profiles on the training set:

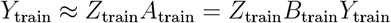

Where 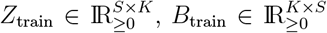. Then, to validate the model on the held-out test set, we use the training archetypes to perform prediction on the test set. More precisely, we use the *A*_test_ = *B*_train_*Y*_test_ as the archetypal features on the test set and keep *Z*_train_ as the corresponding loading matrix.

To tune the regularization hyperparameter, we apply a 3-fold cross-validation (CV) procedure by splitting *Y*_train_ further into 3 independent subsets and using two of them for training and one for validation for every fold. We use Pearson correlation as the scoring metric, and pick the best regularization parameter from 50 values evenly spaced between −7 and 5 on a log scale. Picking the hyperparameter with best mean performance, we use it to retrain the model on *Y*_train_ and test the final performance of the model on *Y*_test_ as described above.

### ArchVelo model

We build an ordinary differential equation (ODE) model of chromatin dynamics, transcription, splicing and degradation cascade for every gene. We propose that gene transcription can be decomposed into a sum of components driven by chromatin dynamics of accessibility archetypes *a*_*k*_(*t*), and assume individual kinetics at each archetype. Following MultiVelo,^15^ we model dynamics of chromatin opening and closing using a simple two-state model. Normalizing each chromatin accessibility archetype between 0 and 1, we assume that it asymptotically approaches 1 in the open state and asymptotically decays to 0 in the closed state: 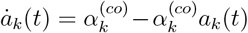 for some rate 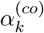 if chromatin is opening and 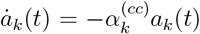 for some rate 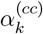 if it is closing. This modeling choice was motivated in MultiVelo by empirical observation and by an assumption that initially rapid chromatin remodeling gets slowed down by biochemical constraints such as the structures of histone complexes and their inter-molecular interactions.^15^ With the simplifying assumption that the chromatin opening and closing kinetics are mirror images of each other, i.e., opening and closing have the same rates, we can join these equations into one: 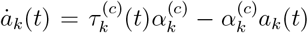

Where

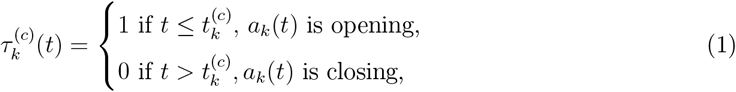

and we assume individual chromatin switch times 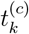 for every archetype. In what follows we will assume 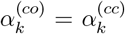 just for simplicity of notation while the ArchVelo model fits these rates separately.

We aim to solve the following system of ODEs:

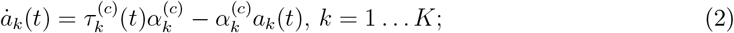

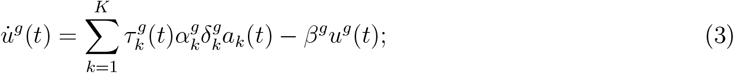

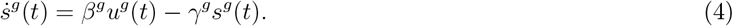

Here eq. (2) describes chromatin dynamics while eqs. (3) and (4) describe dynamics of unspliced and spliced counts, respectively. We assume that the transcription can be active and repressed at every archetype, with 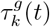 a piecewise constant function analogous to eq. (1) switching from 0 to 1 at transcription initialization time 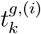 and back to 0 at transcription repression time 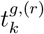. We assume that every archetype has its own transcription kinetic parameters, with every archetype *a*_*k*_, when active, contributing to transcription with a gene-dependent rate 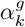. Simultaneously, for every gene *g*, we set transcription rate at an archetype to be proportional to its accessibility score for *g*, i.e., its total loading for the peak summits mapping to 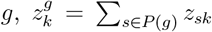. Finally, we correct for prior normalization of the chromatin archetypes here by renormalizing them back by *n*_*k*_ = max(*a*_*k*_) − min(*a*_*k*_). This results in 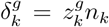 term in eq. (3). Finally, *β*^*g*^ and *γ*^*g*^ denote splicing and degradation rates, respectively.

Due to the linear nature of the above model, *u*^*g*^ and *s*^*g*^ can be decomposed into a linear combination of unspliced and spliced profiles resulting from transcription at every archetype: 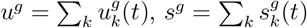 where 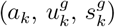 solve a system of ODEs

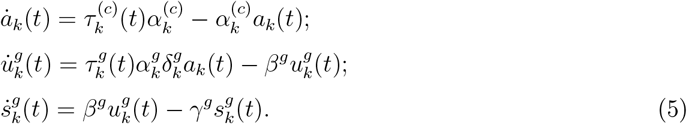

For every *k*, this system mimics a MultiVelo model.^15^ Each system can be solved analytically, with smooth solutions for every continuous interval of time characterized by the same chromatin and transcriptional state. However, we emphasize that the solution to the ArchVelo problem cannot be obtained by the repeated application of MultiVelo at every archetype. The optimization is performed jointly for all *k* due to the presence of shared parameters *β*^*g*^, *γ*^*g*^ and jointly defined likelihood. We iteratively update the model parameters and latent time assignments in an expectation-maximization procedure (EM) which is adapted to the ArchVelo model likelihood (eq. (6) below).

### Parameter initialization for ArchVelo

Due to an increase in the number of parameters in ArchVelo compared to MultiVelo, we sought to find a suitable initialization for ArchVelo parameters. To this end, on the same dataset, we first apply a modification of MultiVelo that we call MultiVelo-AA. MultiVelo-AA differs from MultiVelo in how the chromatin accessibility profiles for individual genes are defined. In MultiVelo, a single chromatin accessibility profile *c*^*g*^ for every gene is defined using the sum of accessibility profiles at its promoter and other linked peaks.^15^ In MultiVelo-AA, the chromatin profiles for every gene are instead predicted using the results of AA on the Pearson normalized ATAC matrix (see **Archetypal analysis on ATAC modality**). More precisely, for every gene *g*, in MultiVelo-AA, we use 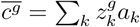 in lieu of the aggregated chromatin accessibility profile *c*^*g*^ and MultiVelo is run on the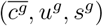 triple. We use the output of MultiVelo-AA to initialize latent time assignments, and kinetic parameters for transcription, splicing and degradation for every gene. More precisely, while the chromatin dynamics parameters are fit separately for every archetype, for other kinetic parameters, the same initialization is used for all components. Note that, in both ArchVelo and MultiVelo-AA, weighted nearest neighbor (WNN) smoothing is first applied to the archetypes *a*_*k*_ according to MultiVelo procedure, yielding a set of smoothed archetypes. These smoothed archetypes are used in ArchVelo to fit eq. (2) and in MultiVelo-AA to define 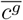; we will nevertheless not include any smoothing notation and will keep denoting the archetypes by *a*_*k*_ in the description of ArchVelo.

### Archetypal velocity components

Velocity profiles 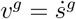 for individual genes are obtained from the solution of eq. (4). The velocity graph (transition matrix for the cells), common latent time assignment and velocity embeddings are produced via the corresponding methods in scVelo package.^13^ By default, as in MultiVelo, to bring the velocity profiles for individual genes to the same scale, we apply *l*_1_ normalization to each of them. By default, likelihood cutoff 0.05 is used in ArchVelo.

Archetypal decomposition of chromatin dynamics automatically provides downstream linear decomposition for the velocity profiles 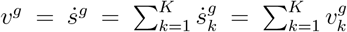 (eq. (5)). As a result, cell trajectories can be decomposed into components potentially driven by different regulatory events. Moreover, the individual latent time estimates for every component can provide a more accurate ordering of cells in case of conflicting signal or presence of cell cycle component. The normalized velocity 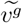 can then be decomposed into a sum of normalized velocity components 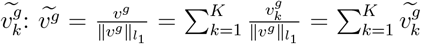. Note that the universal normalization with the *l*_1_ norm of the full velocity is applied to each component, rather than individual *l*_1_ normalization. The velocity graph, latent time assignments and velocity embedding can then be calculated for each *k* to infer archetypal velocity fields. In MultiVelo and ArchVelo, velocity profiles are smoothed among nearest neighbors using the connectivity matrix of the RNA modality; however, the unsmoothed velocities are stored in ArchVelo and can be used.

Archetypal velocity components allow to define cell-specific archetypal velocity loadings that can aid in downstream cell and gene selection. More precisely, we define cell-specific archetypal velocity loadings as 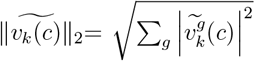 (see **Fig. S2A,C** for an example). Cell-specific archetypal velocity loadings can potentially highlight cells sharing a regulatory program. Furthermore, we also leverage the archetypal decomposition to cluster genes using their spliced count decomposition. Specifically, genes are grouped according to the spliced count component with the highest *l*_1_-norm (see **Fig. S2B,D** for an example). Gene selection also aids in defining the common latent time for an individual component. To calculate it, we min-max normalize the gene-specific latent time estimates and take the average over the genes from the corresponding archetypal cluster.

For archetypal velocity components, we also developed a variation of the velocity embedding visualization on a grid. In scVelo and MultiVelo, the velocity graph is built using the cosine similarity between the velocity vector and the vector of change in gene expression between cells. This approach is agnostic to the cell-specific norm of the velocity vector and is not effective for visualizing the velocity components which are usually concentrated on a subset of the manifold. To alleviate this problem, we developed a modification of the scVelo grid embedding technique which scales the velocity components back by 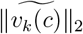 estimates on the grid. This method was applied to visualize all archetypal velocity components in this paper.

### ArchVelo model likelihood

We assume Gaussian noise in the observations which come in the form of spliced, unspliced counts and archetypal chromatin accessibility profiles from AA on the ATAC modality *Y*. The archetypes are weighed with their corresponding weights 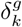 (see **ArchVelo model**). Additionally, we introduce a parameter *w*_*c*_ scaling the influence of *Y* on the velocity inference. Denoting 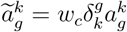, for cell *i*, the observation becomes 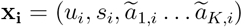. For the prediction of the form 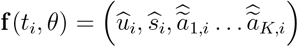, we optimize the negative log-likelihood:

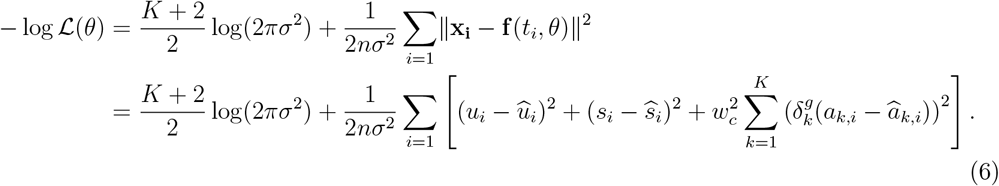

Weight *w*_*c*_ is also available as a parameter in MultiVelo, with *w*_*c*_ = 0.6 at default. For benchmarking experiments, *w*_*c*_ was varied between 0 and 1, with 11 different parameters at increments of 0.1. As the result of benchmarking, for ArchVelo, *w*_*c*_ = 0.3 was chosen as the default parameter.

### ArchVelo expectation-maximization details

We iteratively update the model parameters and latent time assignments in an expectation-maximization procedure (EM) adapted to the model likelihood. More precisely, during the maximization step, the model parameters are updated via maximum likelihood (equivalent to minimizing the mean squared error). For each *k* = 1 … *K*, the model includes 2 + 6*K* parameters: 3*K* parameters describing chromatin dynamics 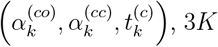 parameters for the transcription component 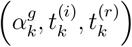, and 2 parameters for splicing and degradation. We first update the parameters fitting eq. (2) to the archetypes using current latent time estimates (via the dual annealing method implemented in scipy Python package). We then optimize the component of the likelihood corresponding to eqs. (3, 4) to update the parameters corresponding to transcription, splicing and degradation. We use the Nelder-Mead method implementation from scipy Python package to optimize the parameters.

The E-step updates the latent time assignment of every cell to its expected value given the current updated ODE parameters. We adopt the strategy described in MultiVelo for the E-step. There the uniformly spaced anchor points are maintained, and cells are assigned with the latent time of the nearest anchor point. 500 anchor points are similarly maintained by default. The key difference here is the nearest neighbor graph which utilizes the distances in the (*K* +2)-dimensional phase space of unspliced, spliced counts and *K* archetypes rather than the 3-dimensional space of (*c, u, s*) in MultiVelo (see **ArchVelo model likelihood**). Nearest neighbor assignment is obtained via the kd-tree quick nearest neighbor lookup implemented in scipy Python package.

### Benchmarking metrics and details

To benchmark ArchVelo against MultiVelo and scVelo, we used two metrics: latent time silhouette and mean log-likelihood in the *u*–*s* phase space. With the underlying model assuming a unique common latent time profile, an appropriate metric of robustness should measure the degree of alignment of latent time estimates. To achieve this, we correlate gene-specific inferred latent time profiles using Spearman correlation. We have found that, typically, individual latent time assignments of scVelo and MultiVelo are correlated between genes within several gene modules and anticorrelated between those modules. To assess latent time robustness, we apply hierarchical clustering to the Spearman correlation profiles (average method, Euclidean metric) and cut the tree to form *n* = 2, …, 20 clusters. For every *n*, we calculate silhouette score (silhouette_score method implemented in sklearn Python package) of the resulting clustering. Mean silhouette score over *n* is reported as latent time silhouette. Simultaneously, for comparison of ArchVelo and MultiVelo, we use their common (*u, s*) component of the log-likelihood to assess the quality of the fit. To obtain a fair likelihood comparison, the same subset of cells was used for both methods for every gene. For this, we used the cell subset chosen by the outlier removal procedure in MultiVelo which could potentially only serve in favor of the MultiVelo method. Finally, we note that, in all benchmarking experiments, for every method, we used default parameters (except *w*_*c*_ which was varied for the benchmark) and the same set of cells and genes and, as a result, the same kNN graph.

To evaluate cross-boundary direction correctness, we used cross_boundary_correctness method from VeloAE Python package, version 0.2.0. For the mouse embryonic brain, we constructed the ground truth graph for the neuronal compartment only. The graph contains two branches originating from IPCs. IPCs have been shown to be capable to produce both upper-layer (UL) and deeper-layer (DL) neurons.^36, 37^ In the UL branch, a pathway to UL neurons from Unc5d^+^ precursors has been previously described.^38, 39^ For the deeper layer, we used *Bcl11b* (*Ctip2*), a well-known marker of DL, for establishing cluster identities.^40^ Expression of *Bcl11b* placed AG.1-4 clusters into the DL branch, and we united them into one cluster which we named AG. Strong axon guidance signature expression and intermediate levels of other well-known markers of DL identity led us to place AG between IPC and DL on this branch. For the HSC dataset, we constructed the ground truth graph for all clusters. Here we refer the reader to several reviews on hematopoiesis.^46, 64, 65^ Previous publications suggested an early split between the neutrophil-monocyte-lymphoid lineage and the BEM-erythroid-megakaryocyte progenitors.^43, 45–47^ The common progenitor to the latter lineage was termed erythro-myeloid progenitor (EMP),^45^ the notation that we adopt. Notably, our ground truth model of hematopoiesis is also similar to the model reconstructed by Zheng et al.^48^ in another single-cell study. Finally, we note that while MEPs are capable of giving rise to bona fide basophils in basophilic conditions,^43^ we included an additional MEP-to-BEM edge to refine the previously available model.

### ArchVelo transition scores

To calculate ArchVelo transition probabilities, cell by cell matrix of transition probabilities was extracted applying transition_matrix method from scvelo.tools on the ArchVelo velocity field. For this analysis, to fix the number of neighbors used for every cell for normalization purposes, the velocity field was recalculated with parameter mode_neighbors = ‘connectivities’. By default, we use 10 nearest neighbors for this analysis. Transition probabilities were then averaged over every pair of clusters (to normalize for cluster sizes) and normalized to condition on the source cluster.

### Single-cell ATAC-seq data preprocessing

All single-cell analysis was done using scanpy.^66^ By default, for all ArchVelo analyses, the summit counts were filtered to the ones present in at least 1% of cells. For every peak summit count matrix *Y*, we applied Poisson correction ⌈*Y/*2⌉ on the raw counts.^67^ Our default normalization step before archetypal analysis was to apply Pearson residual approach to the resulting matrix, with *θ* = 1 as the default overdispersion parameter. The residuals were clipped at 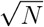 where *N* is the number of cells. We used an implementation of this normalization technique from Python scanpy package.

### Single-cell RNA-seq data preprocessing

Scanpy was used for all analysis of scRNA-seq data. Typical QC steps were followed to remove cells based on low or extremely high signal as well as high mitochondrial counts. For initial analysis, normalization was performed (Pearson residuals or log-transformed library size normalized counts, depending on dataset, see below) and then standard scRNA-seq processing steps were followed: genes with low counts were filtered, highly variable genes were selected, PCA was run, kNN graph was built, and clustering performed. By default, standard imputation step was run for unspliced and spliced counts on each dataset with the same parameters that were used for kNN graph construction. Cell types were determined based on marker gene expression. For differential expression analysis, rank_genes_groups from scanpy with method = ‘wilcoxon’ was used. Full details of RNA-seq analysis run on each dataset can be found below.

### BABEL gene expression predictions

BABEL code was downloaded from github and then edited to enable the assignment of train-test splits to compare directly using the same training and test sets as our archetypal regression. Each dataset was then run using the train_model.py function with mouse v25 gencode for mouse datasets or the HG38 version supplied in the BABEL codebase. Anndatas were read in and modified to be in the expected format for BABEL (i.e., concatenating the ATAC and RNA modalities into one anndata). The RNA predictions from ATAC were extracted from the resulting atac_rna_adata.h5ad anndata. Pearson correlations were calculated with the true gene expression values using np.log1p on the predicted *X* matrix and log library size normalization on the real gene expression values.

### ArchR gene scores

Gene scores were generated using the addGeneScoreMatrix with all default parameters. The gene score matrix was extracted using the ‘getMatrixFromProject’ function, with useMatrix = “GeneScoreMatrix”.

### Peak and peak summit calling

MACS2 was run on single base pair Tn5 insertion sites for all datasets to generate the set of peaks. Additional 150bp-wide summit regions of maximum signal within these peaks were then generated, using bigwig files of signal in the peak regions, using the biosignals package with a custom script. Counts over these 150 basepair regions (referred to as peak summits throughout the text) were generated (typically using ArchR, dataset-specific details can be found below) and used for all subsequent analyses.

### Nearest genes mapping

Gene annotations were used to assign peak summit regions to nearest genes, using the “bedtools closest” function. For mouse datasets, the GTF annotations file from Gencode V25 was used for mouse mm10 genome, downloaded from https://ftp.ebi.ac.uk/pub/databases/gencode/Gencode_mouse/release_M25/. For human, HG38 v46 annotation was downloaded from Gencode at https://ftp.ebi.ac.uk/pub/databases/gencode/Gencode\_human/release\_46.

Peak summits were sequentially labeled as promoter (if within 2kb), exon (if directly within an exon), intronic (if distance to gene was 0), or intergenic (if within 50kb). The rest of the summits were left unlabeled.

### TF motif analysis

Motif binding weights for TF summits were generated using motifmatchr^68^ with a p-value threshold of 1e-3, over the 693 clustered motif PWMs from Vierstra et al.^69^ For the CD8 T cell dataset, motif binding weights were generated using PWMs from a set of TFs focused on T cells from Pritykin et al.^56^ Thus, summit by motif matrix of motif binding weights *M* was generated for each analysis. Cell-specific motif scores were defined as the columns of *Y* ^*T*^ *M*, where *Y* is the summit by cell pearson-normalized matrix of ATAC counts. These scores are shown on UMAPs in **Fig. 5C-E**. Then motif correlation scores *c*_*m,a*_ for every motif *m* and archetype *a* were defined as the Pearson correlation between *a* and the vector of cell-specific scores corresponding to *m*. In **Fig. 7F**, correlation scores were calculated for the progenitor compartment only. We further argued that for full datasets the archetypal analysis results can be also used to calculate the motif scores more directly as *Z*^*T*^ *M* where *Z* is the summit by archetype matrix of archetypal loadings. Thus, in **Fig. 5C-E**, the motifs were first filtered by their archetypal correlation score (*c*_*m,a*_ *>* 0.1) and then the motif score was shown using barplots.

### PBMC data analysis

Data was downloaded from the 10x website.^31^

#### ATAC modality

An initial analysis was performed to determine a final set of cells to use, with ArchR. ArrowFiles were generated using the ‘createArrowFiles’ function, setting minTSS=4, minFrags = 1000, addTileMat=TRUE, and addGeneScoreMat = True. Doublets were inferred using the ‘addDoubletScores’ function, using k=10, knnMethod = “UMAP”, and LSIMethod = 1. An ArchR project was then created using these ArrowFiles, and 1,249 doublets were filtered using the ‘filterDoublets’ function, with a filterRatio set to 1. These cells were then used as the final cells for subsequent analysis.

A bed file of single-base pair insertion sites was generated from the fragment file, only incorporating this final list of cells. Peaks were called using this bed file with MACS2 callpeak, using shift set to −75 and extsize set to 150, qvalue to 0.05, with nomodel, bdg to True, SPMR to True, and keep-dup to all. 150bp-wide summit regions of maximum signal within the peaks were generated using the biosignals package with a custom script.

To generate the peak summit read count matrix, a new set of ArrowFiles was generated, using minTSS=0. minFrags=1, maxFrags=1e20, addTileMat = TRUE, and addGeneScoreMat=TRUE. An ArchRProject was then generated on these ArrowFiles, and then filtered to the cell list generated previously, using the subsetCells function. Next, addIterativeLSI, addClusters, addUMAP, and addImputeWeights were all running using default parameters. The peak summit read count matrix was generated using addPeakSet and addPeakMatrix (run with a ceiling of 20). The peak summit read count matrix was retrieved using the ‘getMatrixFromProject’ function with useMatrix = “PeakMatrix”. This was then read into scanpy, and filtered to peaks present in at least 1% of cells.

#### RNA modality

The filtered feature matrix was downloaded from 10x website and loaded into scanpy. Genes present in fewer than 1% of cells were filtered out (23,118 genes), and then cells with fewer than 1% of this set of genes was filtered out (23 cells). 309 cells with more than 30% mitochondrial reads were removed, 14 cells with total counts greater than 20k were removed, and 26 cells with more than 5000 genes present were removed. Pearson residual normalization was run with theta of 100. PCA was run with 100 PCs. The scRNA-seq matrix was then subsetted to the cells present in the scATAC-seq modality, resulting in 9,952 cells. The KNN graph was built using cosine metric, 30 nearest neighbors, and 30 PCs, and UMAP was run with all default parameters. Leiden clustering was performed with resolution of 1.2, and cell type annotations were determined based on marker gene expression. Specifically, MS4A1, CD19, and CD79A were used to determine B cells; CD3D, CD3E, and CD4 were used to classify CD4 T cells, some of which were assigned as Tregs based on expression of FOXP3 and IKZF2; CD3D, CD3E, CD8A, and CD8B1 were used to identify CD8 T cells, of which some were defined as Activated CD8 if also expressing GZMK, PRF1, and KLRG1, the rest expressed high levels of CCR7, TCF7, and SELL and were assigned as CD8 mem/naïve; a few cells expressed low levels of CD3G, CD4, CD8A, and CD8B1 and were assigned as T cells; NK cells were determined using NKG7 and NCAM1; pDCS were identified using TCF4, MZB1, IL3RA, IRF4, SERPINF1, PLAC8, LILRB4, GZMB, JCHAIN, IRF7, SLC15A4, SPIB, ITM2C, and IRF8; CD14+ monocytes were identified using LYZ, CD14, VCAN, CD36, and SERPIN1A; and CD16+ monocytes were identified using TCF7L2, MTSS1, SERPINA1, RHOC, MS4A7, IFITM2, FCGR3A, SIGLEC10, CX3CR1, and LILRB1.

### Mouse brain data analysis

Data was downloaded from the 10x website.^34^

#### ATAC modality

Single-base pair insertion sites were generated from the fragment file. Peaks were called for each cell type separately based on the original cell type annotations from the paper, using MACS2, with shift set to −75 and extsize set to 150, qvalue to 0.05, bdg to True, SPMR to True, and keep-dup to all. These peaks were then merged using the R package GenomicRanges’ reduce function. A bigwig was generated for each cell type using the bedGraphToBigWig tool. 150 bp summit regions of maximum signal within the peaks were generated using the biosignals package by adding the signal from these bigwigs together in an R script.

The peak summit read count matrix was generated using ArchR. ArrowFiles were first generated from the fragment file, using minTSS=0. minFrags=1, maxFrags=1e20, addTileMat = TRUE, and addGeneScoreMat=TRUE. An ArchRProject was then generated on these ArrowFiles. The peak summit matrix was generated using the addPeakMatrix function with a ceiling of 1000. It was then retrieved using the ‘getMatrixFromProject’function with useMatrix = “PeakMatrix” and binarize set to FALSE. This matrix was then filtered to the cells that were used in MultiVelo. After standard preprocessing steps described above, archetypal analysis was applied with *k* = 10 on this dataset.

#### RNA modality

The cells were first filtered according to the MultiVelo tutorial resulting in 3653 cells. Initially, a liberal cutoff of min_shared_counts=10 via filter_and_normalize method from scVelo package was applied. Log-transformed library normalized counts were used for this dataset. 30 PCs and 50 nearest neighbors were used to construct the kNN graph for this dataset and Leiden clustering with resolution=4 was applied to identify subpopulations relevant to our analysis. Two small subpopulations were filtered out: one based on expression of vascular-associated ECM and adhesion markers including *Col4a1/2, Lama4, Itgb1*, and one based on expression of GABAergic neuron markers such as *Gad2, Nrxn3*, and *Dlx1/2*. The resulting matrix was filtered again with parameters min_shared_counts = 10 and 1000 top highly variable genes were selected. The final matrix after cells and genes were filtered for high-quality ATAC signal contained 3252 cells and 930 genes. The kNN graph for this matrix was recalculated with the same parameters. Cluster annotation was performed according to the available single-cell mouse embryonic brain atlas.^70^ More precisely, for our neuronal compartment, we used the gene signatures resulting from the hierarchical Poisson factorization on all cells in the neuronal compartment of their dataset (see Extended Data Fig. 3d for the atlas^70^). For the glial subpopulations, we used the markers expressed by glioblasts in this atlas (see Extended Data Fig. 6e for the atlas^70^).

### HSC data analysis

Data for day 7 (MV2) was downloaded from GEO using accession number GSE209878.

#### ATAC modality

Single-base pair insertion sites were generated from the fragment file over cells from the paper. The same pipeline as for the mouse brain data was run, with all same parameters. The final peak summit matrix from ArchR was filtered so that all cells contained >3000 total summit count. Summits were filtered according to the standard preprocessing pipeline described above. Archetypal analysis was applied with *k* = 9 to this dataset.

#### RNA modality

14895 cells in the original dataset were first filtered according to standard QC pipelines and filtered for doublets using scrublet algorithm^71^ available in Python through scanpy resulting in 12289 cells. Genes were filtered with parameters min_cells = 20, min_shared_counts=100. To annotate this dataset, due to a strong cell cycle signal, cell cycle regression was performed. We applied the standard pipeline for cell cycle regression, with a slight modification allowing to add the intercept back after regression. The cell cycle regression was performed on the log-transformed library size normalized counts. 1500 highly variable genes were then selected, and 50 PCs and 50 nearest neighbors were used for this dataset. Data was clustered with resolution=3 and re-annotated based on available literature. More precisely, we used the following subpopulation markers. HSC/MPP: *HLF, AVP, HOXA9*, LMPP: *FLT3, TCF4*, Prog DC: *LYZ, IRF8, JAML*, GMP: *MPO*, Prog N: *ELANE, AZU1, PRTN3*, EMP: *GATA2, TFRC, CPA3, RHEX*, MEP: *TFRC, TFR2*, Ery: *KLF1, HBB*, Prog MK: *ITGA2B*, Platelet: *PLEK, PF4*. The final matrix after cells and genes were filtered for high-quality ATAC signal contained 10935 cells and 1325 genes. The kNN graph was recalculated for this matrix using 50 PCs and 50 nearest neighbors.

### CD8 T cell data analysis

The CD8 T cell dataset analyzed here was generated as part of a larger study characterizing splenic T cells in LCMV infection.^54^ We focused on the subset of cells previously annotated as effector CD8 T cell, progenitor CD8 T cell and exhausted CD8 T cell subpopulations.

#### ATAC modality

Archetypal analysis was applied on ATAC data from both clones simultaneously. Our standard preprocessing pipeline was applied to the ATAC matrix, followed by archetypal analysis with *k* = 9.

#### RNA modality

The same pipeline was applied to both clones. Genes were first filtered with filter_and_normalize method from scVelo package with min_shared_counts = 30, n_top_genes = 1500 parameters. Pearson residual normalized counts from the full dataset were then used.^54^ 50 PCs, 50 nearest neighbors and cosine metric were used for kNN graph generation (and unspliced and spliced count imputation). We found that the cosine metric is beneficial compared to the Euclidean metric for analysis of the cell cycle dynamics. For MultiVelo, standard preprocessing was applied, with cells filtered to counts>2000 of aggregated ATAC. The final matrix for velocity analyses (after cells and genes were filtered for high-quality ATAC signal) contained 3707 cells and 1410 genes for the Armstrong clone dataset and 2520 cells and 1439 genes for clone 13. To annotate this dataset, cells from both clones (6409 effector, progenitor and exhausted Cd8 T cells prior to cell filtering) were jointly processed. We performed this analysis on the union of genes for both datasets. For finer clustering, we used 50 PCs, 30 nearest neighbors for the kNN graph and performed Leiden clustering with resolution 2. This yielded 23 clusters that were initially grouped into 10 cell types/states. To further split the progenitor subpopulation into Ccr6^−^ and Ccr6^+^ we applied Leiden clustering to this subpopulation with resolution 0.4. For cell ordering over latent time in the progenitor compartment, we used the difference between archetype 5 and archetype 8. To calculate scores for genes that were previously reported to be differentially expressed in the CD62L^+^ and CD62L^−^ precursor exhausted T cell clusters,^61^ we used score_genes method from scanpy.

## Data and code availability

The open-source ArchVelo package is available at https://github.com/pritykinlab/ArchVelo. The computational code used in the analysis presented in this manuscript is available at https://github.com/pritykinlab/ArchVelo_notebooks.

## Acknowledgments

We thank all members of the Pritykin lab and the Developmental Dynamics group at the Flatiron Institute for helpful discussions. We thank Lucy Reading-Ikkanda (Flatiron Institute) for assistance with the graphic design of figures. The Flatiron Institute is a division of the Simons Foundation. This work was supported by the NIH/NIAID grant DP2AI171161, AACR-Bristol-Myers Squibb Immuno-oncology Research Fellowship (19-40-15-PRIT) and the Ludwig Institute for Cancer Research (Y.P.); Boehringer Ingelheim, the Austrian Science Fund (FWF, 10.55776/PAT4163824), and the ERC grant ERC-2023-STG 101116251 (J.v.d.V); NIH grants P30 CA008748 and R01 AI034206 (A.Y.R.). A.Y.R. is an investigator with the Howard Hughes Medical Institute.

## Supplementary figures

**Figure S1.**
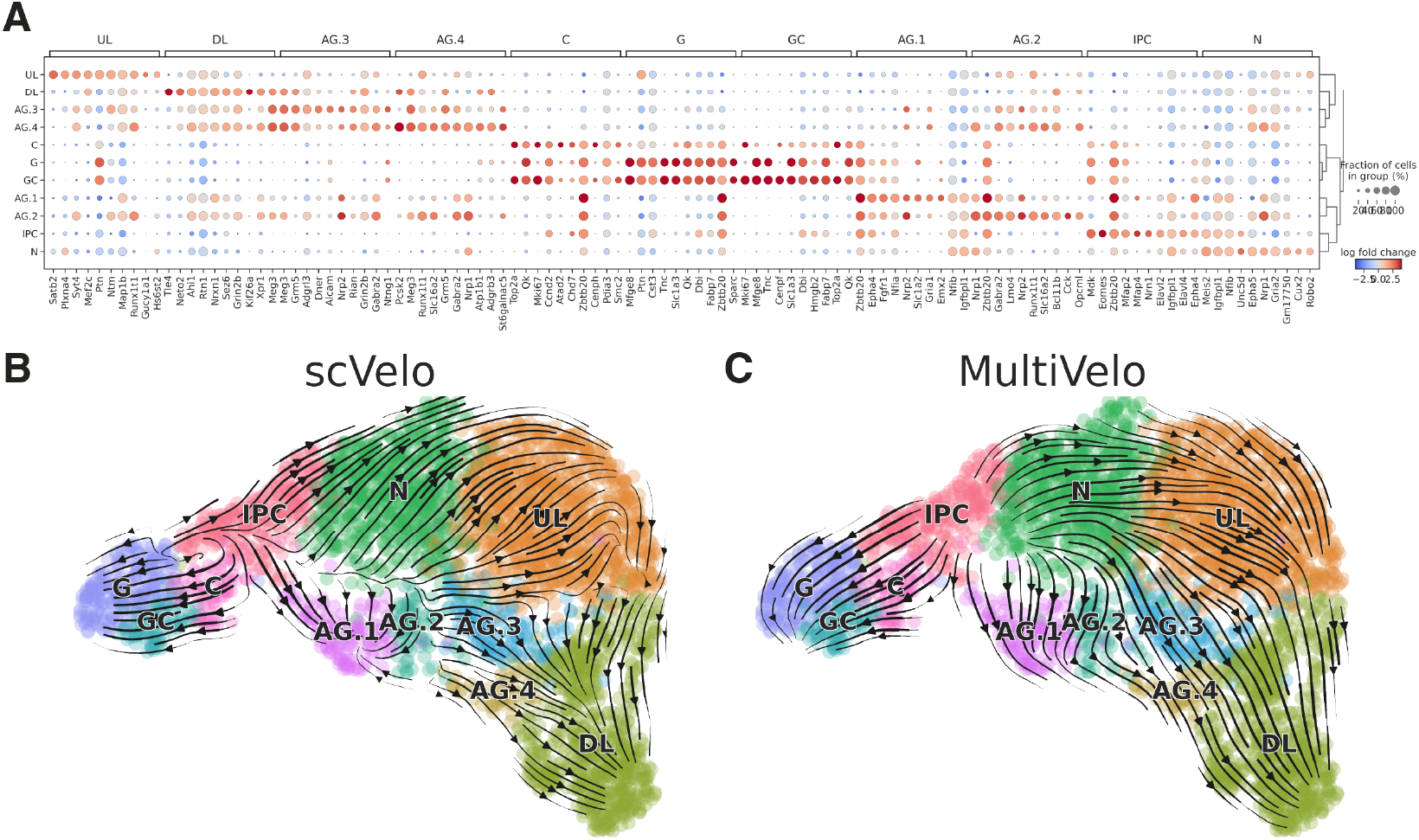
Results of scVelo and MultiVelo on the mouse embryonic brain dataset. **(A)** Differential expression analysis between the identified clusters. For each cluster, the top 10 most significantly differentially expressed genes are shown. Color indicates the log-fold change of mean expression in the cluster compared to all other cells. **(B)** Velocity fields inferred by scVelo, shown on the same UMAP embedding as in **Fig. 2A**. **(C)** Velocity fields inferred by MultiVelo, shown on the same UMAP embedding as in **Fig. 2A**.

**Figure S2.**
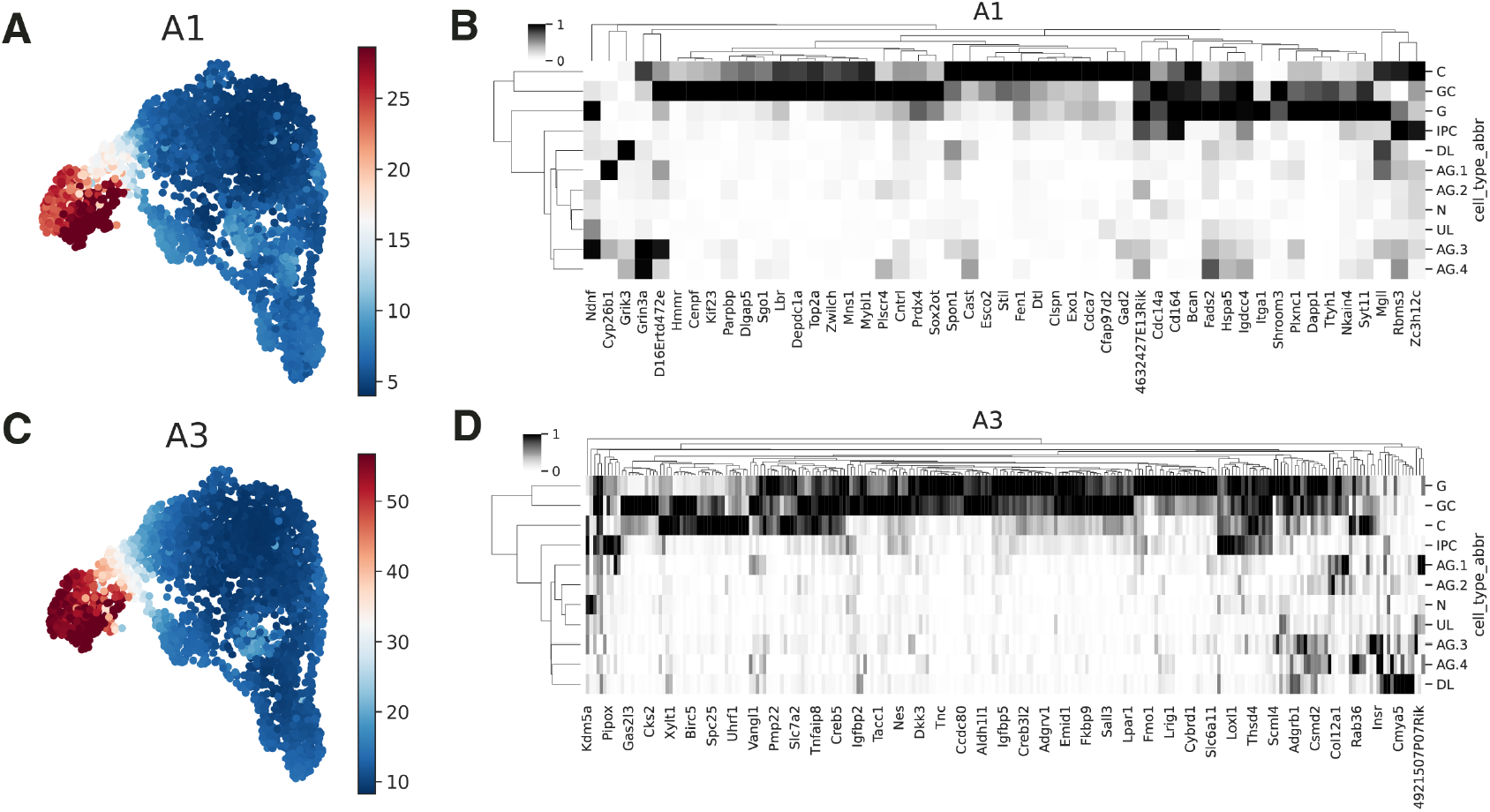
Velocity loadings and gene selection for archetypal components in the mouse embryonic brain dataset. **(A)** UMAP embedding colored by velocity loadings for archetype A1. Loadings are shown as the *l*_2_-norms of the inferred velocity component associated with A1 (see **Methods**). **(B)** Top genes contributing to archetype A1, selected as described in **Methods**. **(C)** Same as panel A, for archetype A3. **(D)** Same as panel B, for archetype A3.

**Figure S3.**
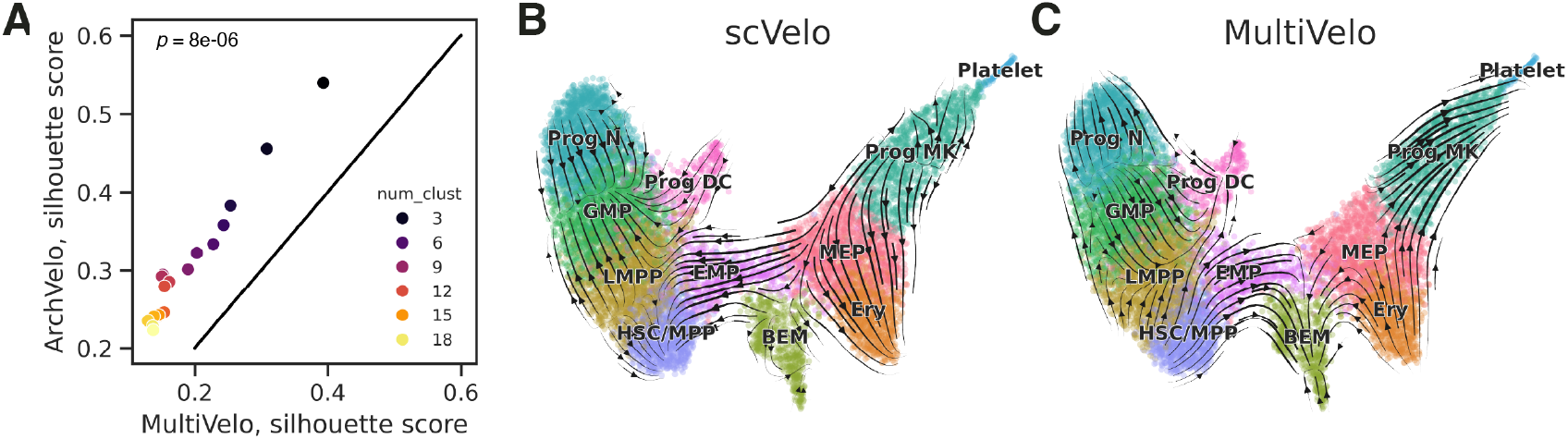
Results of scVelo and MultiVelo on the human hematopoietic stem cell dataset. **(A)** Scatterplot of silhouette scores for latent time correlation clusters (Methods) across varying cluster numbers. *x*-axis: MultiVelo; *y*-axis: ArchVelo. Significance was assessed using the two-sided Wilcoxon signed-rank test. **(B)** UMAP of the HSC differentiation dataset with cell type annotations and velocity fields inferred by scVelo. Cell type annotations are the same as in Fig. 3A. **(C)** Same as panel B, for MultiVelo.

**Figure S4.**
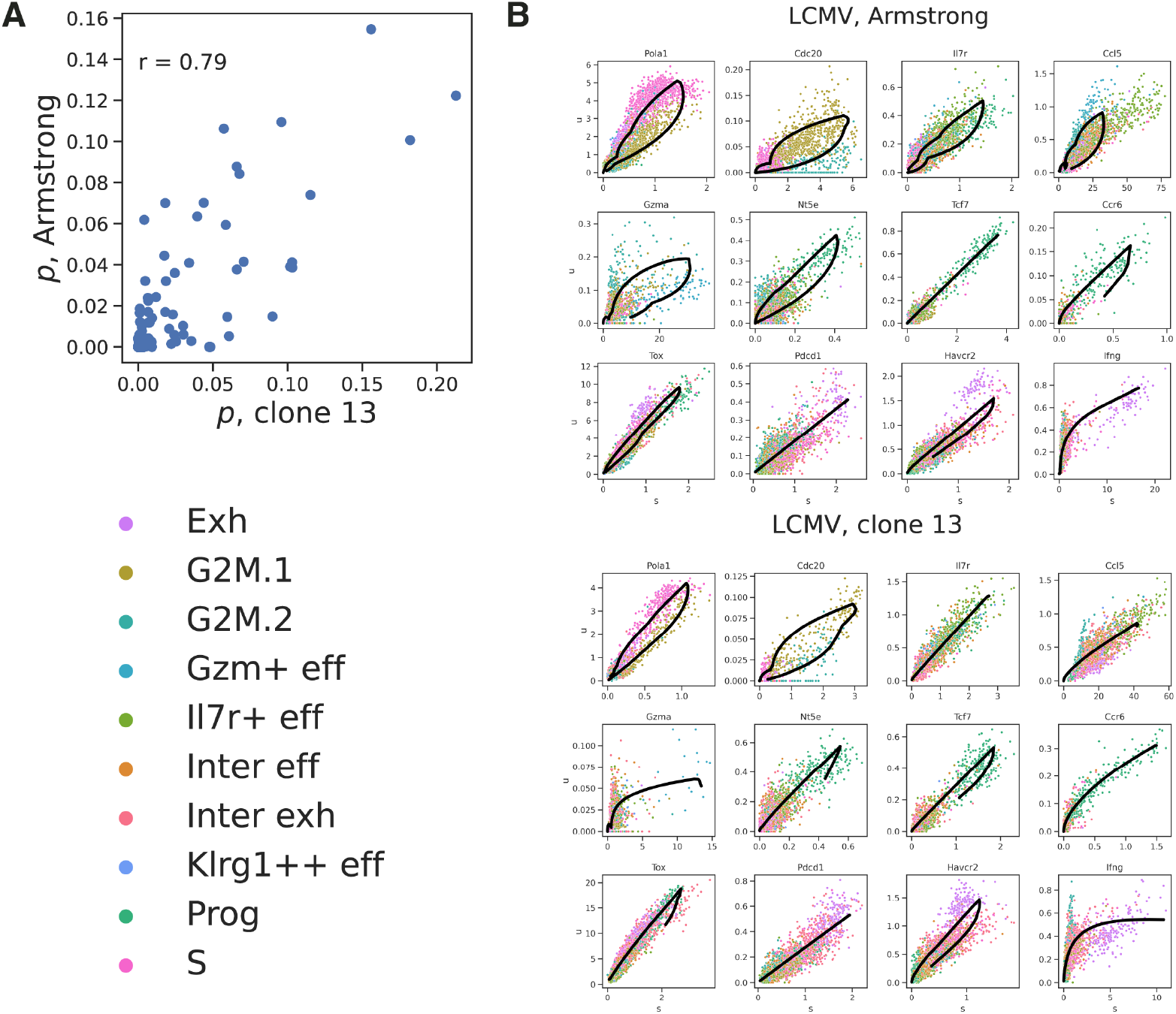
Additional details of the analysis of the CD8 T cell dataset. **(A)** Scatterplot comparing ArchVelo transition probabilities between the two LCMV infection conditions. Each point corresponds to a cluster pair. *x*-axis: clone 13; *y*-axis: Armstrong. Pearson correlation coefficient *r* = 0.79. **(B)** Phase plots for selected genes in LCMV Armstrong and clone 13. Black curve: ArchVelo fit.

**Figure S5.**
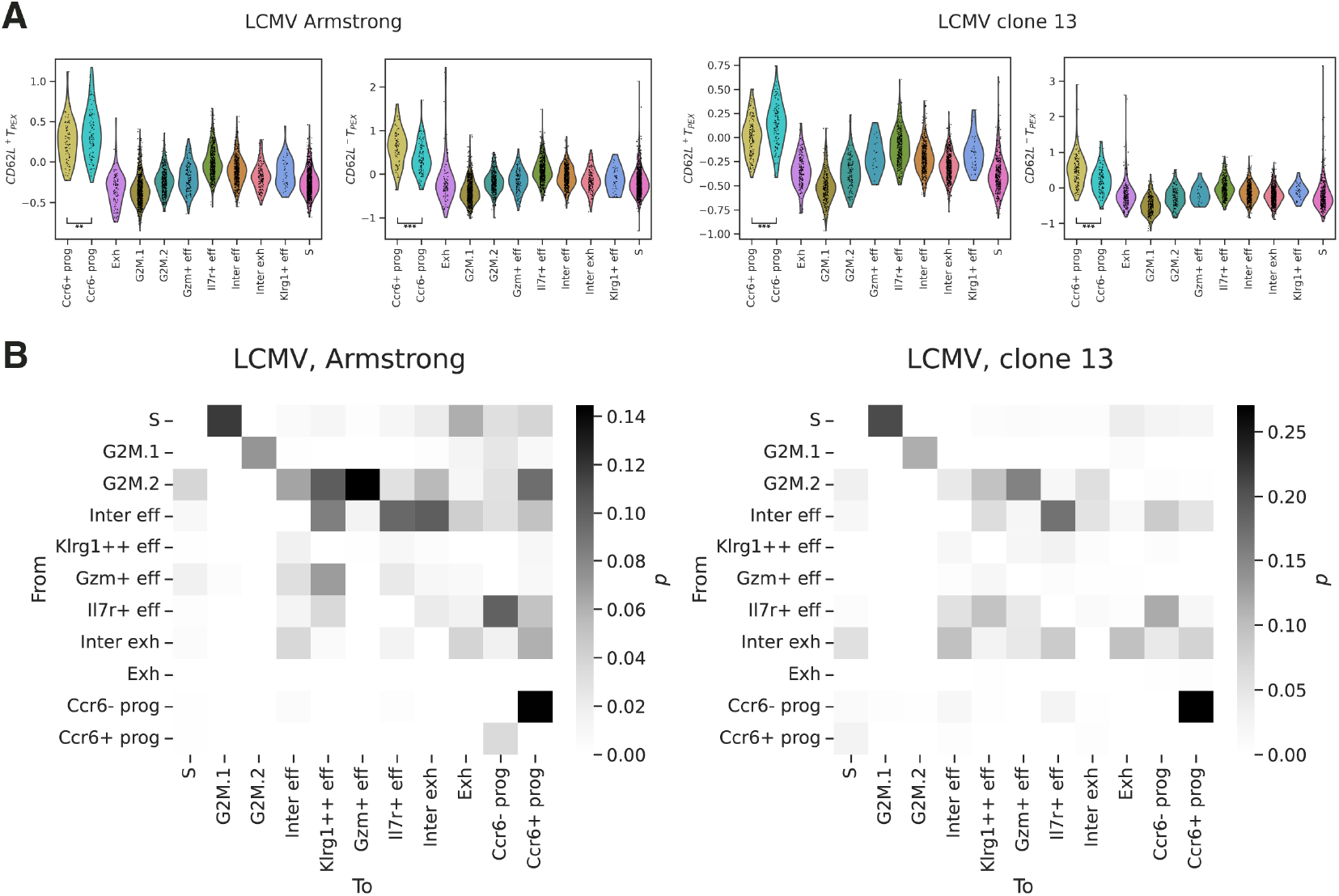
Additional details of the analysis of Ccr6^+^ and Ccr6^−^ CD8 T cell progenitor subsets. **(A)** Violin plots of gene scores for genes differentially expressed in the CD62L^+^ and CD62L^−^ precursor exhausted T (T_PEX_) cell clusters reported in a previous study.^61^ **, *p <* 0.01; ***, *p <* 0.001, one-sided Mann-Whitney U test. **(B)** Heatmaps of ArchVelo-inferred transition probabilities (**Methods**) between clusters including the Ccr6^+^ and Ccr6^−^ CD8 T cell progenitors, shown separately for Armstrong (left) and clone 13 (right).

## References

1 A. Tanay and A. Regev. Scaling single-cell genomics from phenomenology to mechanism. Nature, 541(7637):331–338, 2017.

2 S. Aldridge and S. A. Teichmann. Single cell transcriptomics comes of age. Nature communications, 11(1):4307, 2020.

3 R. Jones, J. Karkanias, M. Krasnow, A. Pisco, S. Quake, J. Salzman, N. Yosef, B. Bulthaup, P. Brown, W. Harper, et al. Tabula Sapiens Consortium*. (2022) The Tabula Sapiens: A multiple-organ, single-cell transcriptomic atlas of humans. Science, 376:.

4 A. Baysoy, Z. Bai, R. Satija, and R. Fan. The technological landscape and applications of single-cell multi-omics. Nature Reviews Molecular Cell Biology, 24(10):695–713, 2023.

5 A. Regev, S. A. Teichmann, E. S. Lander, I. Amit, C. Benoist, E. Birney, B. Bodenmiller, P. Campbell, P. Carninci, M. Clatworthy, et al. The human cell atlas. elife, 6:e27041, 2017.

6 L. Heumos, A. C. Schaar, C. Lance, A. Litinetskaya, F. Drost, L. Zappia, M. D. Lücken, D. C. Strobl, J. Henao, F. Curion, et al. Best practices for single-cell analysis across modalities. Nature Reviews Genetics, 24(8):550–572, 2023.

7 P. V. Kharchenko. The triumphs and limitations of computational methods for scRNA-seq. Nature methods, 18(7):723–732, 2021.

8 M. Efremova and S. A. Teichmann. Computational methods for single-cell omics across modalities. Nature methods, 17(1):14–17, 2020.

9 S. Domcke and J. Shendure. A reference cell tree will serve science better than a reference cell atlas. Cell, 186(6):1103–1114, 2023.

10 D. E. Wagner and A. M. Klein. Lineage tracing meets single-cell omics: opportunities and challenges. Nature Reviews Genetics, 21(7):410–427, 2020.

11 W. Saelens, R. Cannoodt, H. Todorov, and Y. Saeys. A comparison of single-cell trajectory inference methods. Nature biotechnology, 37(5):547–554, 2019.

12 G. La Manno, R. Soldatov, A. Zeisel, E. Braun, H. Hochgerner, V. Petukhov, K. Lidschreiber, M. E. Kastriti, P. Lönnerberg, A. Furlan, et al. RNA velocity of single cells. Nature, 560(7719):494–498, 2018.

13 V. Bergen, M. Lange, S. Peidli, F. A. Wolf, and F. J. Theis. Generalizing RNA velocity to transient cell states through dynamical modeling. Nature biotechnology, 38(12):1408–1414, 2020.

14 V. Bergen, R. A. Soldatov, P. V. Kharchenko, and F. J. Theis. RNA velocity—current challenges and future perspectives. Molecular systems biology, 17(8):e10282, 2021.

15 C. Li, M. C. Virgilio, K. L. Collins, and J. D. Welch. Multi-omic single-cell velocity models epigenome–transcriptome interactions and improves cell fate prediction. Nature biotechnology, 41(3):387–398, 2023.

16 M. Gao, C. Qiao, and Y. Huang. UniTVelo: temporally unified RNA velocity reinforces single-cell trajectory inference. Nature Communications, 13(1):6586, 2022.

17 A. Riba, A. Oravecz, M. Durik, S. Jiménez, V. Alunni, M. Cerciat, M. Jung, C. Keime, W. M. Keyes, and N. Molina. Cell cycle gene regulation dynamics revealed by RNA velocity and deep-learning. Nature communications, 13(1):2865, 2022.

18 M. Lange, V. Bergen, M. Klein, M. Setty, B. Reuter, M. Bakhti, H. Lickert, M. Ansari, J. Schniering, H. B. Schiller, et al. CellRank for directed single-cell fate mapping. Nature methods, 19(2):159–170, 2022.

19 Y. Gu, D. Blaauw, and J. D. Welch. Bayesian inference of rna velocity from multi-lineage single-cell data. bioRxiv, pages 2022–07, 2022.

20 A. Gayoso, P. Weiler, M. Lotfollahi, D. Klein, J. Hong, A. Streets, F. J. Theis, and N. Yosef. Deep generative modeling of transcriptional dynamics for RNA velocity analysis in single cells. Nature methods, 21(1):50–59, 2024.

21 G. Gorin, M. Fang, T. Chari, and L. Pachter. RNA velocity unraveled. PLOS Computational Biology, 18(9):e1010492, 2022.

22 J. D. Buenrostro, B. Wu, U. M. Litzenburger, D. Ruff, M. L. Gonzales, M. P. Snyder, H. Y. Chang, and W. J. Greenleaf. Single-cell chromatin accessibility reveals principles of regulatory variation. Nature, 523(7561):486–490, 2015.

23 D. A. Cusanovich, A. J. Hill, D. Aghamirzaie, R. M. Daza, H. A. Pliner, J. B. Berletch, G. N. Filippova, X. Huang, L. Christiansen, W. S. DeWitt, et al. A single-cell atlas of in vivo mammalian chromatin accessibility. Cell, 174(5):1309–1324, 2018.

24 K. Zhang, J. D. Hocker, M. Miller, X. Hou, J. Chiou, O. B. Poirion, Y. Qiu, Y. E. Li, K. J. Gaulton, A. Wang, et al. A single-cell atlas of chromatin accessibility in the human genome. Cell, 184(24):5985–6001, 2021.

25 S. Preissl, K. J. Gaulton, and B. Ren. Characterizing cis-regulatory elements using single-cell epigenomics. Nature Reviews Genetics, 24(1):21–43, 2023.

26 A. Cutler and L. Breiman. Archetypal Analysis. Technometrics, 36(4):338–347, 1994.

27 T. Hastie, R. Tibshirani, J. H. Friedman, and J. H. Friedman. The elements of statistical learning: data mining, inference, and prediction, volume 2. Springer, 2009.

28 L. M. McLane, M. S. Abdel-Hakeem, and E. J. Wherry. CD8 T Cell Exhaustion During Chronic Viral Infection and Cancer. Annu Rev Immunol, 37:457–495, April 2019.

29 M. Hashimoto, A. O. Kamphorst, S. J. Im, H. T. Kissick, R. N. Pillai, S. S. Ramalingam, K. Araki, and R. Ahmed. CD8 T Cell Exhaustion in Chronic Infection and Cancer: Opportunities for Interventions. Annu Rev Med, 69(69):301–318, January 2018.

30 A. Kallies, D. Zehn, and D. T. Utzschneider. Precursor exhausted T cells: key to successful immunotherapy? Nature Reviews Immunology, 2019. PMID: 31591533.

31 Single Cell Multiome ATAC + Gene Expression dataset analyzed using Cell Ranger ARC 2.0.0. PBMC from a Healthy Donor - Granulocytes Removed Through Cell Sorting (10k). https://www.10xgenomics.com/datasets/pbmc-from-a-healthy-donor-granulocytes-removed-through-cell-sorting-10-k-1-standard-2-0-0. Accessed: 2024-10-20.

32 J. M. Granja, M. R. Corces, S. E. Pierce, S. T. Bagdatli, H. Choudhry, H. Y. Chang, and W. J. Greenleaf. ArchR is a scalable software package for integrative single-cell chromatin accessibility analysis. Nature genetics, 53(3):403–411, 2021.

33 K. E. Wu, K. E. Yost, H. Y. Chang, and J. Zou. BABEL enables cross-modality translation between multiomic profiles at single-cell resolution. Proceedings of the National Academy of Sciences, 118(15):e2023070118, 2021.

34 Single Cell Multiome ATAC + Gene Expression dataset analyzed using Cell Ranger ARC 1.0.0. Fresh Embryonic E18 Mouse Brain (5k). https://www.10xgenomics.com/datasets/fresh-embryonic-e-18-mouse-brain-5-k-1-standard-1-0-0. Accessed: 2024-10-20.

35 S. Linnarsson and S. A. Teichmann. Single-cell genomics: coming of age. Genome Biology, 17(1):97, 2016.

36 N. A. Vasistha, F. García-Moreno, S. Arora, A. F. P. Cheung, S. J. Arnold, E. J. Robertson, and Z. Molnár. Cortical and Clonal Contribution of Tbr2 Expressing Progenitors in the Developing Mouse Brain. Cerebral cortex (New York, N.Y.: 1991), 25(10):3290–3302, oct 2015. PMID: 24927931.

37 T. Kowalczyk, A. Pontious, C. Englund, R. A. M. Daza, F. Bedogni, R. Hodge, A. Attardo, C. Bell, W. B. Huttner, and R. F. Hevner. Intermediate neuronal progenitors (basal progenitors) produce pyramidal-projection neurons for all layers of cerebral cortex. Cerebral cortex (New York, N.Y. : 1991), 19(10):2439–2450, oct 2009. PMID: 19168665.

38 V. Tarabykin, A. Stoykova, N. Usman, and P. Gruss. Cortical upper layer neurons derive from the subventricular zone as indicated by Svet1 gene expression. Development (Cambridge, England), 128(11):1983–1993, jun 2001. PMID: 11493521.

39 K. Toma and C. Hanashima. Switching modes in corticogenesis: mechanisms of neuronal subtype transitions and integration in the cerebral cortex. Frontiers in neuroscience, 9:274, 2015. PMID: 26321900.

40 B. J. Molyneaux, P. Arlotta, J. R. L. Menezes, and J. D. Macklis. Neuronal subtype specification in the cerebral cortex. Nature Reviews Neuroscience, 8(6):427–437, 2007.

41 C. Qiao and Y. Huang. Representation learning of RNA velocity reveals robust cell transitions. Proceedings of the National Academy of Sciences of the United States of America, 118(49), dec 2021. PMID: 34873054.

42 A. M. Ranzoni, A. Tangherloni, I. Berest, S. G. Riva, B. Myers, P. M. Strzelecka, J. Xu, E. Panada, I. Mohorianu, J. B. Zaugg, and A. Cvejic. Integrative Single-Cell RNA-Seq and ATAC-Seq Analysis of Human Developmental Hematopoiesis. Cell stem cell, 28(3):472–487.e7, mar 2021. PMID: 33352111.

43 D. Pellin, M. Loperfido, C. Baricordi, S. L. Wolock, A. Montepeloso, O. K. Weinberg, A. Biffi, A. M. Klein, and L. Biasco. A comprehensive single cell transcriptional landscape of human hematopoietic progenitors. Nature Communications, 10(1):2395, 2019.

44 J. D. Buenrostro, M. R. Corces, C. A. Lareau, B. Wu, A. N. Schep, M. J. Aryee, R. Majeti, H. Y. Chang, and W. J. Greenleaf. Integrated Single-Cell Analysis Maps the Continuous Regulatory Landscape of Human Hematopoietic Differentiation. Cell, 173(6):1535–1548.e16, may 2018. PMID: 29706549.

45 A. Görgens, S. Radtke, M. Möllmann, M. Cross, J. Dürig, P. A. Horn, and B. Giebel. Revision of the Human Hematopoietic Tree: Granulocyte Subtypes Derive from Distinct Hematopoietic Lineages. Cell Reports, 3(5):1539–1552, 2013.

46 N. Schippel and S. Sharma. Dynamics of human hematopoietic stem and progenitor cell differentiation to the erythroid lineage. Experimental Hematology, 123:1–17, jul 2023.

47 R. Drissen, S. Thongjuea, K. Theilgaard-Mönch, and C. Nerlov. Identification of two distinct pathways of human myelopoiesis. Science immunology, 4(35), may 2019. PMID: 31126997.

48 S. Zheng, E. Papalexi, A. Butler, W. Stephenson, and R. Satija. Molecular transitions in early progenitors during human cord blood hematopoiesis. Molecular Systems Biology, 14(3):e8041. mar 2018.

49 H. Li, P. Côté, M. Kuoch, J. Ezike, K. Frenis, A. Afanassiev, L. Greenstreet, M. Tanaka-Yano, G. Tarantino, S. Zhang, J. Whangbo, V. L. Butty, E. Moiso, M. Falchetti, K. Lu, G. G. Connelly, V. Morris, D. Wang, A. F. Chen, G. Bianchi, G. Q. Daley, S. Garg, D. Liu, S. T. Chou, A. Regev, E. Lummertz da Rocha, G. Schiebinger, and R. G. Rowe. The dynamics of hematopoiesis over the human lifespan. Nature methods, 22(2):422–434, feb 2025. PMID: 39639169.

50 Y. Ben-David, B. Gajendran, K. M. Sample, and E. Zacksenhaus. Current insights into the role of Fli-1 in hematopoiesis and malignant transformation. Cellular and molecular life sciences: CMLS, 79(3):163, feb 2022. PMID: 35412146.

51 L. S. Bastian, B. A. Kwiatkowski, J. Breininger, S. Danner, and G. Roth. Regulation of the megakaryocytic glycoprotein IX promoter by the oncogenic Ets transcription factor Fli-1. Blood, 93(8):2637–2644, apr 1999. PMID: 10194443.

52 Y. Li, X. Qi, B. Liu, and H. Huang. The STAT5–GATA2 pathway is critical in basophil and mast cell differentiation and maintenance. The Journal of Immunology, 194(9):4328–4338, 2015.

53 A. B. Cantor and S. H. Orkin. Transcriptional regulation of erythropoiesis: an affair involving multiple partners. Oncogene, 21(21):3368–3376, may 2002. PMID: 12032775.

54 S. K. Walker, J. van der Veeken, A. Rudensky, and Y. Pritykin. Single-cell multiomics reveals archetypal regulatory programs shared across CD4 and CD8 T cell subsets in viral infection. bioRxiv, 2025.09.08.675014, 2025.

55 Y. Zhong, S. K. Walker, Y. Pritykin, C. S. Leslie, A. Y. Rudensky, and J. van der Veeken. Hierarchical regulation of the resting and activated T cell epigenome by major transcription factor families. Nature immunology, 23:122–134, 2022.

56 Y. Pritykin, J. van der Veeken, A. R. Pine, Y. Zhong, M. Sahin, L. Mazutis, D. Pe’er, A. Y. Rudensky, and C. S. Leslie. A unified atlas of CD8 T cell dysfunctional states in cancer and infection. Molecular Cell, 81(11):2477–2493, 2021.

57 T. Chu, M. Wu, B. Hoellbacher, G. P. de Almeida, C. Wurmser, J. Berner, L. V. Donhauser, A.-K. Gerullis, S. Lin, J. D. Cepeda-Mayorga, I. I. Kilb, L. Bongers, F. Toppeta, P. Strobl, B. Youngblood, A. M. Schulz, A. Zippelius, P. A. Knolle, M. Heinig, C.-P. Hackstein, and D. Zehn. Precursors of exhausted T cells are pre-emptively formed in acute infection. Nature, 640(8059):782–792, April 2025.

58 D. T. McManus, R. M. Valanparambil, C. B. Medina, C. D. Scharer, D. J. McGuire, E. Sobierajska, Y. Hu, D. Y. Chang, A. Wieland, J. Lee, T. H. Nasti, M. Hashimoto, J. L. Ross, N. Prokhnevska, M. A. Cardenas, A. L. Gill, E. C. Clark, K. Abadie, A. J. Kumar, J. Kaye, B. B. Au-Yeung, H. Y. Kueh, H. T. Kissick, and R. Ahmed. An early precursor CD8+ T cell that adapts to acute or chronic viral infection. Nature, 640(8059):772–781, April 2025.

59 C. Gago da Graça, A. A. Sheikh, D. M. Newman, L. Wen, S. Li, J. Shen, Y. Zhang, S. S. Gabriel, D. Chisanga, J. Seow, A. Poch, L. Rausch, M.-H. T. Nguyen, J. Singh, C.-H. Su, L. A. Cluse, C. Tsui, T. N. Burn, S. L. Park, B. Von Scheidt, L. K. Mackay, A. Vasanthakumar, D. Bending, W. Shi, W. Cui, J. Schröder, R. W. Johnstone, A. Kallies, and D. T. Utzschneider. Stem-like memory and precursors of exhausted T cells share a common progenitor defined by ID3 expression. Science Immunology, 10(103):eadn1945. January 2025.

60 M. L. Burger, A. M. Cruz, G. E. Crossland, G. Gaglia, C. C. Ritch, S. E. Blatt, A. Bhutkar, D. Canner, T. Kienka, S. Z. Tavana, A. L. Barandiaran, A. Garmilla, J. M. Schenkel, M. Hillman, I. de Los Rios Kobara, A. Li, A. M. Jaeger, W. L. Hwang, P. M. K. Westcott, M. P. Manos, M. M. Holovatska, F. S. Hodi, A. Regev, S. Santagata, and T. Jacks. Antigen dominance hierarchies shape TCF1(+) progenitor CD8 T cell phenotypes in tumors. Cell, 184(19):4996–5014.e26, sep 2021. PMID: 34534464.

61 C. Tsui, L. Kretschmer, S. Rapelius, S. S. Gabriel, D. Chisanga, K. Knöpper, D. T. Utzschneider, S. Nüssing, Y. Liao, T. Mason, S. V. Torres, S. A. Wilcox, K. Kanev, S. Jarosch, J. Leube, S. L. Nutt, D. Zehn, I. A. Parish, W. Kastenmüller, W. Shi, V. R. Buchholz, and A. Kallies. MYB orchestrates T cell exhaustion and response to checkpoint inhibition. Nature, 609(7926):354–360, 2022.

62 T. J. Stewart and S. I. Abrams. Altered immune function during long-term host-tumor interactions can be modulated to retard autochthonous neoplastic growth. The Journal of Immunology, 179(5):2851–2859, 2007.

63 M. Mørup and L. K. Hansen. Archetypal analysis for machine learning and data mining. Neurocomputing, 80:54–63, 2012.

64 E. Laurenti and B. Göttgens. From haematopoietic stem cells to complex differentiation landscapes. Nature, 553(7689):418–426, 2018.

65 L. A. Liggett and V. G. Sankaran. Unraveling Hematopoiesis through the Lens of Genomics. Cell, 182(6):1384–1400, sep 2020.

66 F. A. Wolf, P. Angerer, and F. J. Theis. SCANPY: large-scale single-cell gene expression data analysis. Genome biology, 19:1–5, 2018.

67 L. D. Martens, D. S. Fischer, V. A. Yépez, F. J. Theis, and J. Gagneur. Modeling fragment counts improves single-cell ATAC-seq analysis. Nature Methods, 21(1):28–31, 2024.

68 A. Schep and S. University. motifmatchr: Fast Motif Matching in R, 2022.

69 J. Vierstra, J. Lazar, R. Sandstrom, J. Halow, K. Lee, D. Bates, M. Diegel, D. Dunn, F. Neri, E. Haugen, E. Rynes, A. Reynolds, J. Nelson, A. Johnson, M. Frerker, M. Buckley, R. Kaul, W. Meuleman, and J. A. Stamatoyannopoulos. Global reference mapping of human transcription factor footprints. Nature, 583(7818):729–736, July 2020.

70 G. La Manno, K. Siletti, A. Furlan, D. Gyllborg, E. Vinsland, A. Mossi Albiach, C. Mattsson Langseth, I. Khven, A. R. Lederer, L. M. Dratva, et al. Molecular architecture of the developing mouse brain. Nature, 596(7870):92–96, 2021.

71 S. L. Wolock, R. Lopez, and A. M. Klein. Scrublet: Computational Identification of Cell Doublets in Single-Cell Transcriptomic Data. Cell Systems, 8(4):281–291.e9, April 2019.

